# Landscape of Large-Scale Somatic Genomic Insertions in Non-Small Cell Lung Carcinoma Revealed by Nanopore Sequencing

**DOI:** 10.1101/2023.10.07.561149

**Authors:** Dan Xie, Lin Xia, Zhoufeng Wang, Tianfu Zeng, Xuenan Pi, Huan Wang, Guonian Zhu, Xinyue Wu, Yangqian Li, Yan Deng, Yawen Qi, Xuyan liu, Fengmei Zhang, Weimin Li

**Affiliations:** Department of Respiratory and Critical Care Medicine, Laboratory of Omics Technology and Bioinformatics, Frontiers Science Center for Disease-related Molecular Network, State Key Laboratory of Biotherapy, West China Hospital, Sichuan University, Chengdu, Sichuan, 610041, China; Institute of Respiratory Health, Frontiers Science Center for Disease-related Molecular Network, West China Hospital, Sichuan University, Chengdu, Sichuan, 610041, China; Precision Medicine Key Laboratory of Sichuan Province, West China Hospital, Sichuan University, Chengdu, Sichuan, 610041, China

**Author notes:** These authors contributed equally (Lin Xia, Zhoufeng Wang, Tianfu Zeng, Xuenan Pi, Huan Wang).

## Abstract

Previous NSCLC genomic studies were mostly based on the next-generation sequencing of short reads, which is an efficient approach for identifying single nucleotide variants and small indels but ineffective for identifying structural variants, especially large-scale insertions. Here, we studied 151 lung adenocarcinoma (LUAD) and 106 lung squamous cell carcinoma (LUSC) samples and paired blood samples using nanopore sequencing technology. We developed a rigorous computational pipeline and characterized the landscape of large-scale somatic insertions in NSCLC. Combining other omics data, we report three findings: 1. we identified an LUSC-enriched somatic simple repeat expansion shared by approximately 40% of LUSC patients that regulates *PTPRZ1* gene expression through distal enhancers; 2. the somatic insertion of transposable elements (TEs) in NSCLC were mostly ‘complex TEs’ consisting of multiple TE elements; and 3. the insertion of short interspersed nuclear elements, especially from the Alu family in young lineages, is a frequent somatic mutation type that shapes the transcriptome of NSCLC through the expression of these elements.

## Introduction

Non-small cell lung carcinoma (NSCLC) is the leading cause of cancer-related death worldwide, and the number of NSCLC patients reported annually is growing rapidly [1]. The presence of somatic mutations within tumor cells is a defining characteristic of tumorigenesis and progression in NSCLC [2–5]. In the past decade, high-frequency somatic mutations in lung carcinoma have been identified by next-generation sequencing (NGS), among which driver mutations, such as mutations in *EGFR*, *KRAS*, and *TP53* [6], have been demonstrated to be clinically relevant [3]. In addition to single nucleotide variants (SNVs) and short insertions and deletions (indels), structural variants (SVs) in the NSCLC genome, such as deletions, translocations, copy-number alterations, and complex genomic rearrangements, have also been explored. The findings include the loss of *ALK* [2], *MET* exon 14 skipping [7], amplification of *KRAS*, and gene fusions involving *ROS1* or *RET* associated with translocations [8] or complex genomic rearrangements [5]. However, few studies have explored large-scale insertions (> 50 bp) in the NSCLC genome due to the limitations of short-read NGS data.

Most previous studies on the NSCLC genome were based on NGS short reads, which are efficient and accurate for identifying SNVs and small indels. However, it is challenging to reliably identify insertions using short reads given that the reads originating from insertions cannot be unambiguously aligned to the genome. Long-read sequencing (LRS) technology, represented by nanopore sequencing technology [9] and PacBio technology [10], can generate reads with lengths of several kilobases. These reads can completely span insertions, thereby enhancing the accuracy of insertion identification. With the rapid development of long-read sequencing technology, several studies have sequenced the genomes of lung cancer cells and identified SVs that have been missed by NGS. Sakamoto et al. sequenced lung cancer cells and a small number of clinical samples on the nanopore sequencing platform and successfully detected a previously unknown complicated aberration pattern referred to as “cancerous local copy-number lesions” (CLCLs) [11]. In addition, using the nanopore sequencing platform, Suzuki et al. revealed precise junctions of gene fusions, such as *EML4-ALK*, in lung adenocarcinoma (LUAD) cells [12]. However, these studies only sequenced a limited number of samples or cell lines, and each focused on only a small number of genomic SVs. None of these studies discussed somatic insertions in primary NSCLC samples.

Large-scale insertions are a predominant type of SV that can be classified in three subtypes: tandem duplications (TDs), transposable elements (TEs) and *de novo* insertions [13, 14]. Somatic TDs may define subtypes of cancer, such as acute myeloid leukemia [15], breast cancer [16] and prostate cancers [17]. A previous study reported that TD-targeting enhancers near *MYC* and *AR* may influence post-translational gene regulation in metastatic prostate cancer [18]. However, the landscape and potential functions of somatic TDs in the NSCLC genome remain enigmatic. Moreover, short-read sequencing data (SRS) do not allow the accurate estimation of TD size, especially for TDs involving short repeat expansions (SREs), even when the most recent bioinformatics tools are applied [19]. TEs play fundamental roles in early embryonic development [20], evolution [21], human diseases [22], and carcinogenesis [23]. Recently, a pancancer study reported the global profile of TE expression [24] and demonstrated the sufficiency of AluJb expression for activating the oncogene LIN28B in lung cancer cell lines. Another study analyzed the transcriptome of LUSC samples from The Cancer Genome Atlas (TCGA) and identified LINE-1-FGGY as the most frequent somatic TE in the Chinese cohort; this TE could serve as a potential prognostic biomarker and therapeutic target for lung squamous cell carcinoma (LUSC) [25]. Moreover, the prevalence of complex INS structures formed through TE recombination has been identified in human diseases, contributing significantly to genomic diversity [26]. Although several studies have reported a limited number and type of TEs in cancer [27, 28], the diversity and complexity of somatic TEs in the context of cancer genomes, including the NSCLC genome, have not been fully characterized.

Here, we sequenced the primary tumor samples of 242 NSCLC patients using nanopore sequencing technology and developed bioinformatics pipelines to reliably identify somatic insertions (INSs) in these tumors. We classified somatic INSs as TDs, TEs or *de novo* insertions (Methods). We identified an LUSC-enriched cluster based on a shared somatic SRE located between distal enhancers that regulated the *PTPRZ1* gene. Data from the independent cohort showed that 39.62% of LUSC but only 1.32% of LUAD samples harbored this somatic SRE. We found that somatic TEs formed from a complex combination of multiple TE segments are highly common in the NSCLC genome, comprising 56.10% of all somatic TEs. Moreover, we identified and verified the widespread expression of somatic TEs in NSCLC, and *Alu* elements dominated the expressed types.

## Results

### Nanopore Sequencing Identifies Large-Scale Insertions in Hundreds of Lung Carcinoma Patients

We collected tumor samples and paired blood samples from 242 stage I-IV NSCLC patients, including 147 LUAD and 95 LUSC patients. All tumors were treatment naive, surgically resected and histologically confirmed to be LUAD or LUSC (the demographic characteristics and pathological findings of the cases are described in Table S1). LUSC was classified as keratinized or nonkeratinized squamous (Figure S1A). A total of 7 multiple synchronous tumors were obtained from 3 of these patients with LUAD (Table S1). As a result, we collected 257 tumor samples and 253 paired blood or normal lung samples. All patients provided written informed consent to participate in genomic studies in accordance with local institutional review boards.

We sequenced the whole genomes of 244 samples (101 LUAD samples, 23 LUSC samples, and 120 paired blood samples) with a PromethION nanopore sequencer and generated long sequencing reads (Table S2). In total, 17.29 terabases (Tb) of long clean reads were generated with an average of 89.97 gigabases (Gb) per tumor sample and an average of 51.01 Gb per paired blood sample (Table S2A). The mean read depth reached 28.5X (16.9X∼49.2X) for the tumor samples and 16X (4.6X∼44.1X) for the paired blood samples. The mean N50 of clean long reads was 24.8 kb (3.79 kb∼44.93 kb). We also sequenced the genomes of 226 of the 244 samples on the Illumina NGS platform with an average coverage of 34.67X (21.84X∼86.52X) for tumor samples and 32.5X (17.13X∼48.24X) for blood samples (Table S2B). In addition, in 84 out of the 124 tumor samples, RNA-sequencing (RNA-seq) datasets were generated on the NGS platform (Table S2C). An assay for transposase-accessible chromatin sequencing (ATAC-seq) [29] dataset was also generated on the NGS platform for 78 of the 124 tumor samples (Table S2D).

Based on the short-read WGS data, we detected 1,712,158 somatic SNVs and indels. The mean somatic mutation rate in our cohort was 4.82 mutations per megabase (Mb) of DNA (median 1.23 Mb), which is lower than the values obtained in the TCGA LUAD and LUSC cohorts [8, 30]. The relative contributions of mutation signatures were calculated by refitting COSMIC consensus mutation signatures by using Mutalisk (Lee et al., 2018), and 8 signatures were reported among our cases (Table S2E). By clustering the mutational signatures, cases were divided into two groups (Figure S1B). The vast majority of cases heavy-smoking harbored a higher proportion of S4 (related to exposure to smoking carcinogens) and a lower proportion of S5 (related to age), which was consistent with their clinical history. Other cases with available smoking history harbored a high proportion of S5. MutSig2CV identified 26 significantly mutated genes (q value < 0.05, Table S2F) in our cohort, and the most prominent cancer-related mutated gene was *EGFR* (50/87 LUAD, 1/22 LUSC), followed by *TP53* (26/87 LUAD, 14/22 LUSC) (Figure S1C). Several cases harbored mutations in known lung cancer driver genes, such as *ERBB2* (11.8%), *KRAS* (7.2%) and *CDKN2A* (4.5%) (Table S2G). Additionally, GISTIC identified focal *EGFR*, *SLC12A7*, and *LETM2* amplifications and focal *APC*, *FAM66A*, and *SPAG11B* deletions, as reported by TCGA (Figure S1D).

To identify high-confidence somatic INSs in each patient using the long sequencing reads, we developed an in-house bioinformatics pipeline (Methods). First, we mapped the long sequencing reads onto the hg38 reference genome using NGMLR software [31]. The average mapping rate was 96.1% (71.13%∼98.93%), indicating high sequencing quality. Second, we applied Sniffles software [31] to call large-scale INSs (>50 bp) for each sample. For each INS locus identified in the tumor samples, we compared the reads from the tumor sample with the reads from the matched blood sample. A candidate somatic INS was identified if multiple reads from the tumor sample supported the INS but no read from the control sample supported the INS. Third, we designed a statistical model-based method to accurately determine the breakpoints of INSs. Fourth, high-confidence insertion loci were confirmed via manual inspection. We assembled and polished each INS locus using Shasta [32], Racon [33] and Medaka [34]. Finally, we characterized distinct classes of INSs based on their assembled inserted sequences. RepeatMasker [35], Tandem Repeat Finder [35] and Sdust [36] were utilized to annotate the assembled inserted sequences. Somatic INSs were classified as TDs, TEs or *de novo* insertions (Methods).

A total of 11,159 somatic INSs with lengths above 50 bp were identified. Notably, no correlation between tumor purity and the number of somatic INSs was observed (ABSOLUTE: mean 0.476; CNV Facets: mean 0.408; Table S2H and S2I, Figure S1E, S1F). To ascertain the fidelity of our somatic INS dataset, we first employed an ONT platform-based ‘Panel of Normals (PONs)’. We developed this panel by pooling structural variants identified from 405 unrelated Chinese individuals [14] and structural variants identified from 63 healthy individuals obtained from the 1000 Genomes Project set [37] together (Methods). All of the WGS data used to construct the PONs were generated using the ONT platform. Only 3 (0.027%) somatic INSs exhibited similarity to the PONs based on criteria such as sequence similarity > 90%, normal allele frequency > 10%, and insertion distance below 100 bp. These 3 low-confidence somatic INSs consisted of 2 TDs and 1 *de novo* INS, all containing simple repeat sequences. Despite differences in the tissue samples and the methods and parameters of sequencing and SV detection used to identify SVs, these results provide further evidence supporting the identified somatic INSs. Next, we estimated the relationship between SV numbers and sequencing depth. We defined eligible regions, which were genomic regions that could be reliably callable in LRS, as 500 bp genomic windows that are captured in at least 90% of blood samples (Methods). A total of 98.78% of somatic INSs fell within eligible regions. These results provided strong evidence of the accuracy and reliability of our somatic INS dataset.

Somatic TDs were the most frequently observed somatic INS type (49.46%), followed by somatic TEs (40.99%) (Figure 1A). We clustered the samples based on the breakpoints of all somatic INSs. The results showed that different samples from the same patient presented high consistency, and the heterogeneity between patients was high (Figure S1G). In addition, we identified a cluster that consisted of LUSC samples. Interestingly, four of the samples, all of which were LUAD samples, exhibited exceptionally higher numbers of somatic INSs (1674∼1902) than the remainder of the samples (2∼261), indicating a subclass with a high INS burden (Figure 1B). Two of the four patients were initially diagnosed with positive metastatic lymph nodes (N2) and developed bone metastasis or brain metastasis within one year after the operation. The other two patients were diagnosed as lymph node-negative (N0) but also experienced recurrence within one year (Table S1). Therefore, the four patients had a poor prognosis, which may be related to the high frequency of somatic INSs.

**Figure 1.**
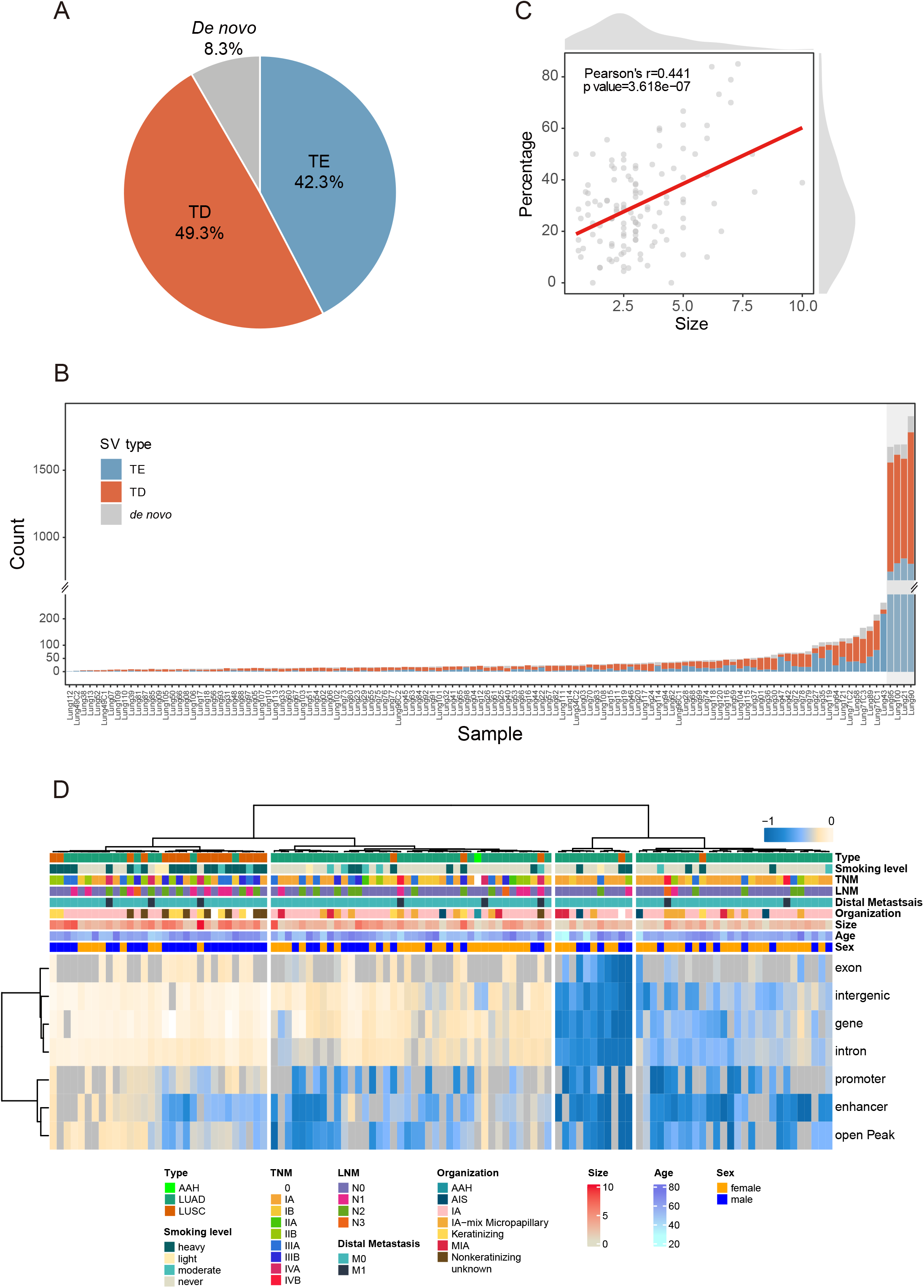
Somatic insertion load investigation. A. Pie chart showing the proportions of detected somatic insertion subtypes (TE: transposable element, TD: tandem duplication, *de novo*: *de novo* insertion). B. The numbers of somatic INS subtypes in each sample are shown (TE: transposable element, TD: tandem duplication, *de novo*: *de novo* insertion). The subclass with a high INS burden is highlighted (gray shaded area). C. Dotplot of the correlation between tumor size and the proportion of somatic TEs among INSs. The density plots show the distribution of tumor size (top) and the proportion of somatic TEs (right). D. Heatmap showing the hierarchical clustering of NSCLC samples based on the Spearman correlation coefficient between the distribution of somatic INSs and SNVs/indels in functional genomic regions. Color indicates the Spearman correlation coefficient range, as in the legend, and gray represents missing values. The open peaks were called using our ATAC-seq data (Methods).

The INS subtype composition was correlated with several clinical parameters. For example, the proportion of somatic TEs among INSs was positively correlated with tumor size (Pearson’s correlation, r = 0.441, p value = 3.618e-07, Figure 1C). Early-stage patients (stage I NSCLC) had a higher proportion of somatic TDs (Wilcoxon rank-sum test, p value = 0.0246) than late-stage patients. These correlations suggested potential functional roles of somatic INSs in the biology of NSCLC.

To explore whether somatic INSs and SNVs/indels tend to affect the same genomic elements, we performed correlation analysis between the distribution of the somatic INS breakpoints and SNVs/indels in functional genomic regions. Surprisingly, the genomic locations of somatic INS breakpoints are generally anti-correlated with those of SNVs/indels, albeit with a weak correlation between the total number of INSs and total number of SNVs/indels (Spearman’s rank correlation, p value < 0.05, correlation score 0.18 for INSs and SNVs, correlation score 0.27 for INSs and indels). We clustered the samples based on correlation scores (Figure 1D) and identified 4 clusters. Cluster 1 had more somatic INSs and SNPs/indels than the other samples (Wilcoxon rank-sum test, p value < 0.01). Cluster 2 displayed a moderate frequency of somatic mutations. Clusters 3 and 4 had fewer somatic SNPs/indels compared to other clusters (Wilcoxon rank-sum test, p value < 1e-05). As SNV/INDEL counts show a negative correlation with INS counts in clusters 3 and 4, we concentrated on the anti-correlation in clusters 1 and 2. We observed that clusters 1 and 2 showed a higher anti-correlation at regulatory regions than at coding elements. Interestingly, most of the LUSC samples were assigned to cluster 1 and presented a similar pattern of anti-correlation with cluster 2. And majority of LUAD samples in clusters 1 had near 0 correlations for coding elements. In addition, patients in cluster 1 tended to have tumors classified in the later tumor-node-metastasis (TNM) stages (chi-square test, p value = 0.00245), have larger tumors than other patients (Wilcoxon rank-sum test, p value = 5.08e-04) and be smokers (chi-square test, p value = 1.25e-05).

### An LUSC-Enriched Somatic SRE Regulates *PTPRZ1* Gene Expression

As shown above (Figure S1G), the clustering of breakpoints of all INS types identified a cluster of LUSC samples. To further explore the details, we investigated the presence of recurrent somatic TDs, TEs or *de novo* INSs in LUSC samples and LUAD samples separately using fishHook [38] (Methods). As expected, we identified 4 recurrent TDs, one of which was shared by 13 LUSC samples (FDR = 8.3e-52, Figure 2A), and the other three were each shared by 3 LUSC samples (Figure 2B). No recurrent somatic INS was identified in LUAD samples. The TD shared by 13 LUSC samples was a somatic simple repeat expansion (SRE; expansion of loci composed of simple repeated motifs) [39] located at chr7: 121603431, a simple repeat region annotated by RepeatMasker. Only 2 LUAD samples shared this somatic SRE. The region originally consisted of a short tandem repeat with a repeat pattern of (TTTC), and the somatic alteration consisted of an expansion of this pattern (Figure 2C). The length of the somatic SRE ranged from 172 to 545 bp (median = 308 bp) among the samples. To validate the presence of somatic SRE in tumors, we performed targeted amplifocation analysis of SRE in tumors and normal tissues of 4 patients. Gel electrophoresis results showed that the DNA fragments from SRE-harboring tumor samples were longer than those from matched blood samples (Figure S2A). A capillary electrophoresis diagram also showed a single peak pattern for the DNA fragments from blood samples, while the SRE-containing tumor samples exhibited a bimodal pattern. These findings confirmed that the somatic SRE existed in LUSC tumor cells (Figure S2B).

**Figure 2.**
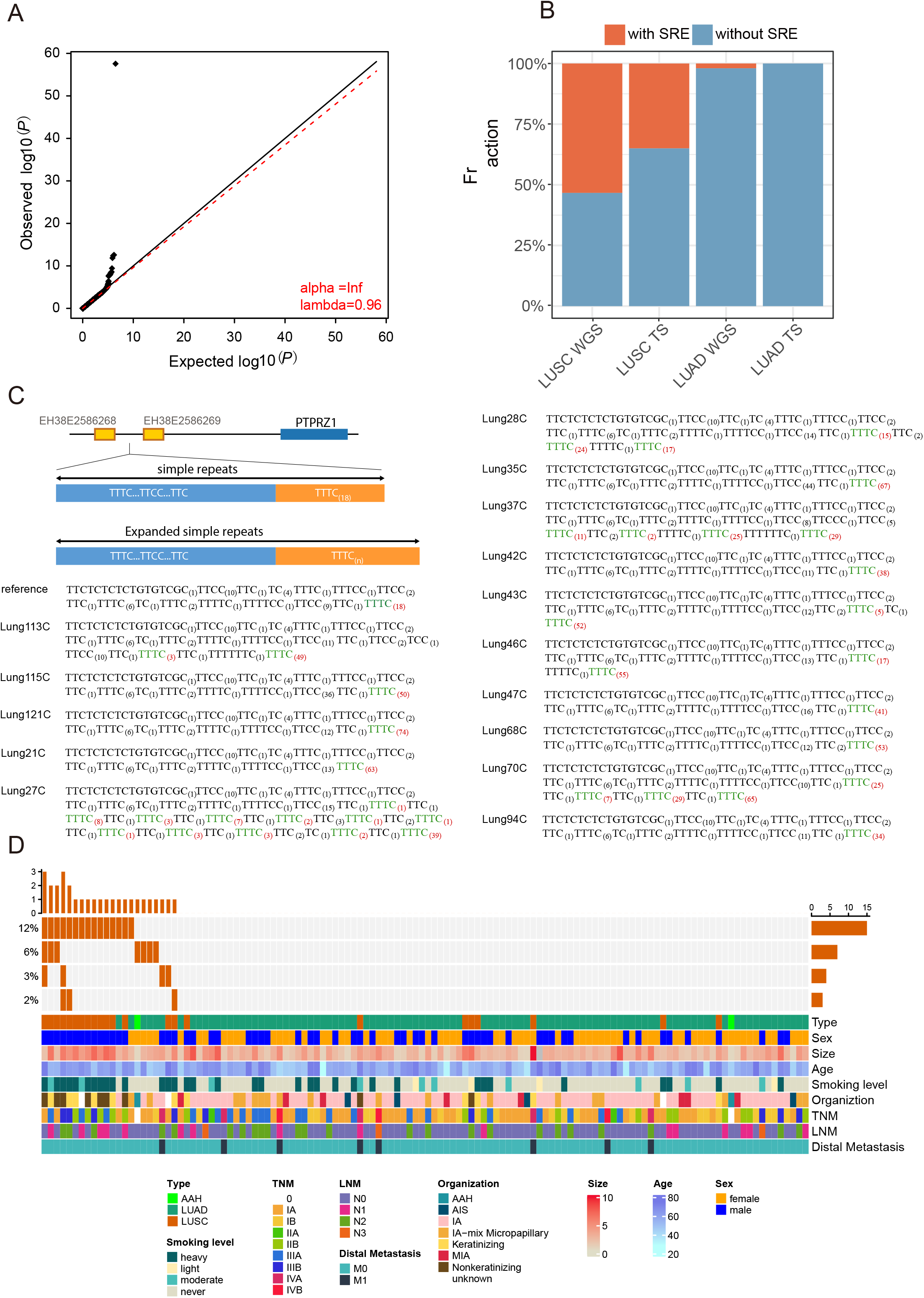
A recurrent somatic SRE may be a marker of LUSC. A. Logarithmic quantile‒quantile (QQ) plot showing the significant enrichment of TDs in LUSC samples reported by fishHook. B. Bar plot showing the fraction of samples with (orange) or without (blue) the shared TD in NSCLC groups (WGS, whole-genome sequencing using the long-read sequencing platform; TS, targeted sequencing using the long-read sequencing platform). C. Schematic representation of the simple repeats located at chr7: 121603431 and the expanded version. The assembled SRE sequences of the 15 samples are shown. The TTTC motif with a variable number of repeats is highlighted in green, and the repeat number is highlighted in red. D. Significantly recurrent TDs (FDR < 0.05 and shared by at least 3 samples) in LUSC samples identified by fishHook are ordered vertically by decreasing prevalence. The percentage of NSCLC samples that had a significantly recurrent TD is noted at the left. Samples are displayed in columns, with the overall number of TDs plotted at the top. TDs are displayed in rows, with the overall sample number plotted at the right.

Intriguingly, this somatic SRE seemed to be enriched in LUSC samples. To validate this hypothesis, we sequenced the whole genomes of an additional 22 pairs of LUSC tumor samples and their matched blood samples on the PromethION platform, and 10 LUSC tumor samples were found to harbor this somatic SRE (Figure 2C). We further sequenced 61 LUSC tumor samples and 50 LUAD tumor samples with matched blood samples targeting the somatic SRE region with a QNome-3841 nanopore sequencer (Figure 2D). Confirming this finding, the somatic SRE was identified in 19 of the LUSC samples but not in the LUAD samples. In total, 39.62% of the LUSC samples sequenced in this study harbored the somatic SRE. In contrast, somatic SREs were noted in less than 1.32% of the LUAD samples.

To ensure that the identified somatic SRE was not a sequencing artifact of LRS, we examined its occurrence in 115 tumor samples and matched blood samples whose WGS SRS data were generated in our study and analyzed with ExpansionHunter Denovo (Dolzhenko et al., 2020) (STAR Method). We found evidence of the existence of the SRE in 14 out of 22 LUSC tumor samples and not in LUAD samples or blood samples (Figure S2C). We then obtained LUSC WGS SRS data from the ICGC LUSC-KR project [40] and found evidence of the existence of the SRE in 16 out of 30 LUSC samples but not in their paired blood samples.

The LUSC-enriched somatic SRE was located in an intergenic region, so we investigated the possible target gene of this mutation. The SRE site was surrounded by two distal enhancers predicted from *SK-N-MC* and *A673* cells [41] (Figure 3A). The target gene of these two distal enhancers has not been reported, but *PTPRZ1*, *FAM3C* and *WNT16* are the nearest protein-coding genes. Published Hi-C data from human embryonic stem cells [42] indicated that several genes, including *PTPRZ1* (Figure 3A), and the two enhancers are located within the same topologically associated domain (TAD) region. A study from the four-dimensional (4D) nucleosome consortium [43] predicted contact between the enhancers and the *PTPRZ1* gene in 3D space, thus suggesting a high possibility of chromatin interaction between the enhancers and the *PTPRZ1* gene (Figure S3A). We experimentally validated the spatial interaction between the enhancers and *PTPRZ1* in primary tumor samples and LUSC cell lines. First, we performed Hi-C on one of the matched clinical samples containing the somatic SRE. The results showed that the enhancer regions, the somatic SRE and *PTPRZ1* gene were located within the same TAD only in the tumor sample and not in the blood sample (Figure 3B). Second, we performed a chromatin conformation capture polymerase chain reaction (3C-PCR) experiment in the LUSC cell line HCC-95, which contained this somatic SRE (Figure S3B), and confirmed the spatial proximity of the SRE-inserted enhancer regions to the *PTPRZ1* gene (Figure S3C-S3E). In contrast, the LUSC cell line H226, which does not contain this somatic SRE, did not exhibit similar spatial proximity between the enhancers and the *PTPRZ1* gene.

**Figure 3.**
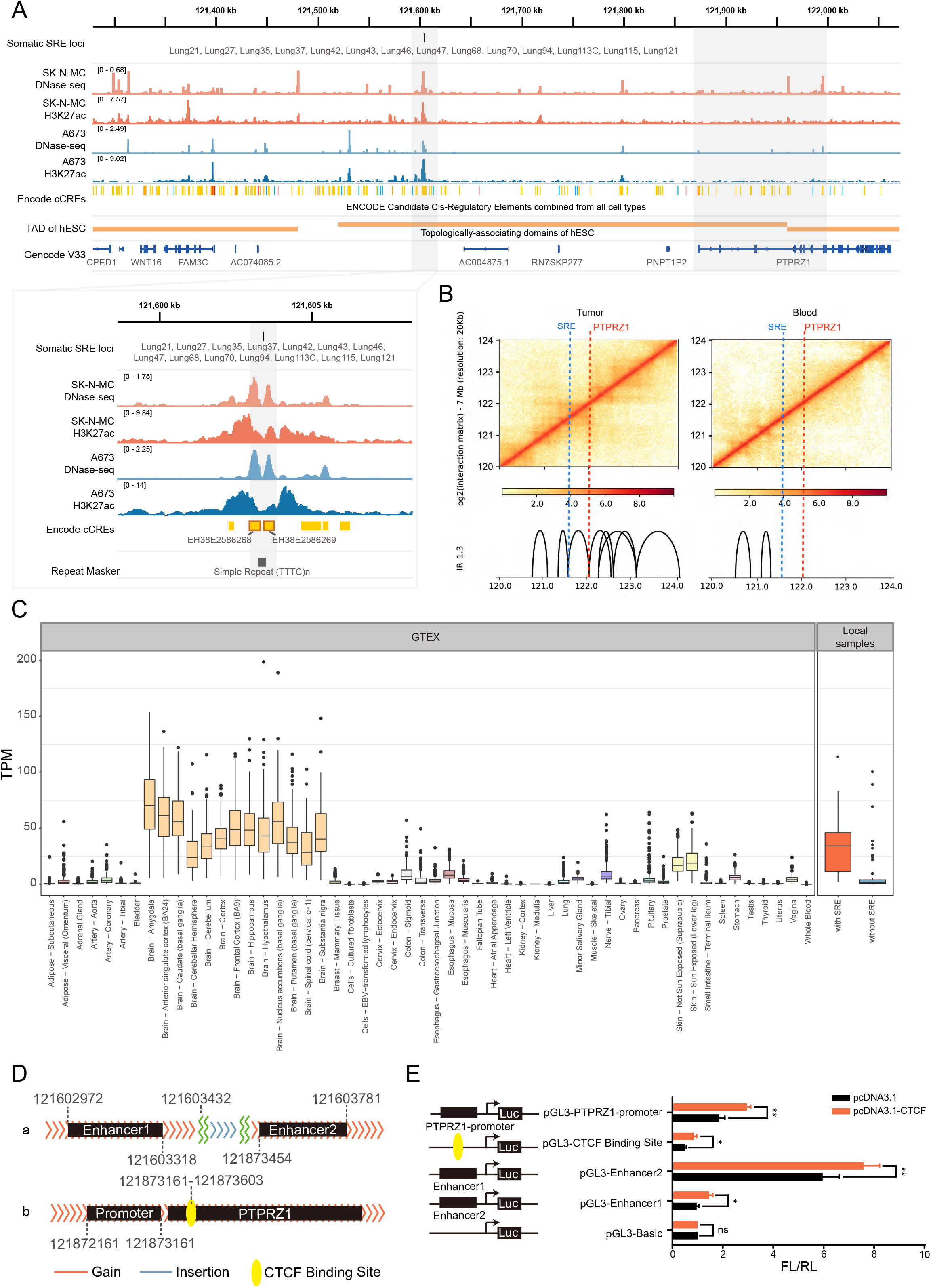
The target of the LUSC-enriched SRE effect may be *PTPRZ1*. A. Browser plot showing the epigenomic signal and TAD around the LUSC-enriched SRE. The enhancers and the simple repeat region (the first gray shaded area) around the LUSC-enriched SRE are shown in the lower left enlarged panel. The overlapping region of *PTPRZ1* and the boundary of the TAD, which harbored the LUSC-enriched SRE, is highlighted (the second gray shaded area). B. The Hi-C interaction matrices around the LUSC-enriched SRE and *PTPRZ1* (20 kb resolution). The left matrix showing the contact map of an LUSC sample, and the right matrix showing the contact map of the matched blood sample of this LUSC sample. The matrices were Knight-RUlz (KR) normalized. In the bottom plot, TADs were scored by an inclusion ratio (IR), which represents the ratio of intra-TAD interactions relative to interactions in regions outside the TAD. TADs with IR > 1.3 are shown with lines. C. Boxplots show the expression level (transcript per million, TPM) of *PTPRZ1* in normal tissues (downloaded from GTEX), NSCLC samples with the LUSC-enriched SRE, and NSCLC samples without the LUSC-enriched SRE. D. Schematic of the possible interaction fragment genomic loci. Panels a and b show the locations of the enhancers (black rectangle), the promoter of *PTPRZ1* (black rectangle), the *PTPRZ1* gene (black rectangle) and the CTCF binding site (yellow oval). E. The dual-luciferase reporter plasmids were constructed by cloning the enhancer1, enhancer2, and *PTPRZ1* promoters and validating the CTCF binding site in the pGL3 plasmid, and CTCF was cloned into the pcDNA3.1 plasmid. HEK293T cells were cotransfected with the indicated vectors and the control or enhancer 1, enhancer 2 or *PTPRZ1* promoter; the Renilla luciferase plasmid (pRL-TK), and pcDNA3.1-CTCF or the control plasmid for 48 hours and then subjected to the luciferase assay. The x-axis is the ratio of firefly luciferase (FL) to Renilla luciferase (RL). *** p value < 0.001, ** p value < 0.01, * p value < 0.05, ns no significant difference.

We next examined whether the somatic SRE affects the *PTPRZ1* expression level by affecting distal enhancers. We tested the differential expression of genes within 500 kb upstream/downstream of the enhancers between tumor samples with and without the somatic SRE. Only *PTPRZ1* showed significantly (Wilcoxon rank-sum test, p value = 3.92e-5, log2|fold change| = 2.34) differential expression. *PTPRZ1* encodes a protein tyrosine phosphatase receptor that is upregulated in the central nervous system [44] and is reported to be related to gliomas. Notably, in our lung carcinoma data, *PTPRZ1* expression levels in samples with the somatic SRE were significantly higher than those in samples without the SRE (Figure 3C). SRE-inserted *PTPRZ1* expression levels were almost as high as those observed in brain tissues (median transcript per million (TPM) = 34.17, Figure 3C). Therefore, it is highly likely that the somatic SRE could affect the expression of *PTPRZ1* through the distal enhancers on either side of the SRE. To validate the regulatory relationship between the distal enhancer and the *PTPRZ1* gene, we performed luciferase reporter assays using HEK293T cells. The dual-luciferase reporter plasmids were constructed by cloning the enhancer, *PTPRZ1* promoter and validated CTCF binding region (Figure 3D) into the pGL3 plasmid, and CTCF was cloned into the pcDNA3.1 plasmid. As per the dual-luciferase reporter system shown in Figure 3E, compared with their respective controls, cotransfection of the *PTPRZ1* reporter, enhancer 1, and enhancer 2 plasmids, separately, with pcDNA3.1-CTCF led to a significant increase in luciferase activity. This result suggested that CTCF could bind to these two enhancers as well as the promoter of *PTPRZ1*. These findings also indicated that potential loops were anchored between these two enhancers and the promoter of *PTPRZ1*, which was mediated by CTCF. Insulator proteins appear indispensable in defining topologically associated domain boundaries and maintaining chromatin loop structures within these domains [45, 46]. We chose to focus on loops anchored by CTCF-binding sites because CTCF plays a well-characterized role in chromatin looping [47–51]. Our results indicated that the two CTCF-binding sites in the enhancer and *PTPRZ1* promoter regions form a boundary insulator element, blocking remote enhancers located within one domain from aberrantly activating promoters in the neighboring domain. Thus, these sites might indirectly ensure the activation of *PTPRZ1* promoters by distal enhancers within the same topological domain.

### Highly expressed PTPRZ1 could promote the proliferation and migration of LUSC tumor cells

Given that the SRE was detected in 39.62% (42/106) of LUSC samples but only 1.32% (2/151) of LUAD samples, we hypothesized that this SRE preferentially occurred in LUSC patients, leading to the differential expression of PTPRZ1 between LUSC and LUAD. We assessed TCGA PTPRZ1 expression data and confirmed that PTPRZ1 was differentially expressed between LUSC and LUAD (adjusted p value = 6.74e-162, log2|fold change| = 3.36, Figure S4A). In addition to central nervous system cancers, skin cutaneous melanoma (SKCM), LUSC and head and neck squamous cell carcinoma (HNSC) presented high PTPRZ1 expression values.

Our results suggested that SRE could distally regulate PTPRZ1 gene expression. To further test this hypothesis, we analyzed the genotype and expression of PTPRZ1 in NSCLC cell lines. Target sequencing results showed that four LUSC cell lines (H520, KNS62, H1703, and HCC95) harbored the SRE, whereas three LUSC cell lines (H226, SK-MES-1 and CALU-1), two LUAD cell lines (A549 and PC-9) and large cell lung cancer H1581 cells did not harbor the SRE (Figure 4A and Table S3). Coincidently, the H520, KNS62, H1703, and HCC95 cell line presented the SRE with length of 73-112 bp (Figure 4B, Table S3). We then investigated PTPRZ1 expression in cell lines with and without the SRE. The RT‒PCR results revealed that the H520 cell line exhibited the highest expression of the PTPRZ1 gene among the examined cell lines. The other four SRE-containing LUSC cell lines (KNS62, H1703 and HCC95) and U251 cells exhibited relatively low expression levels of PTPRZ1, whereas the A549, PC-9 and H1581 and normal bronchus epithelial cells (BEAS-2B) expressed lower levels of PTPRZ1 (Figure 4C). Western blotting was employed to confirm the high expression level of the PTPRZ1 protein in three LUSC cell lines harboring the SRE (Figure 4D). The immunofluorescence assay showed that PTPRZ1 was highly expressed in the cell membrane and cytoplasm of H520 cells and HCC95 cells compared to U251 cells and BEAS-2B cells (Figure 4E-4F). These results implied that the PTPRZ1 expression level was correlated with the presence of the SRE at the cellular level. We then knocked out the SRE in HCC95 cells using the CRISPR/Cas9 gene editing system to investigate whether the SRE affected the migration and invasion ability of these cells, we designed two plasmids containing two guide RNAs targeting the SRE region (Figure S4B). Sanger sequencing showed that a segment of the knockout SRE sequence was lost compared with the reference DNA sequence (Figure S4C). The results of gel electrophoresis and RT-PCR also validated a band of knockout sequence around 300bp in the HCC95 SRE-KO group (Figure S4D), which further indicated that the SRE had been knocked out. And knockout of SRE down regulated the transcription expression of PTPRZ1 (Figure S4E).The cell function results showed that SRE knockout significantly inhibited cell migration ability compared with the control and NC groups in HCC95 cells (Figure S4F-S4I).

**Figure 4.**
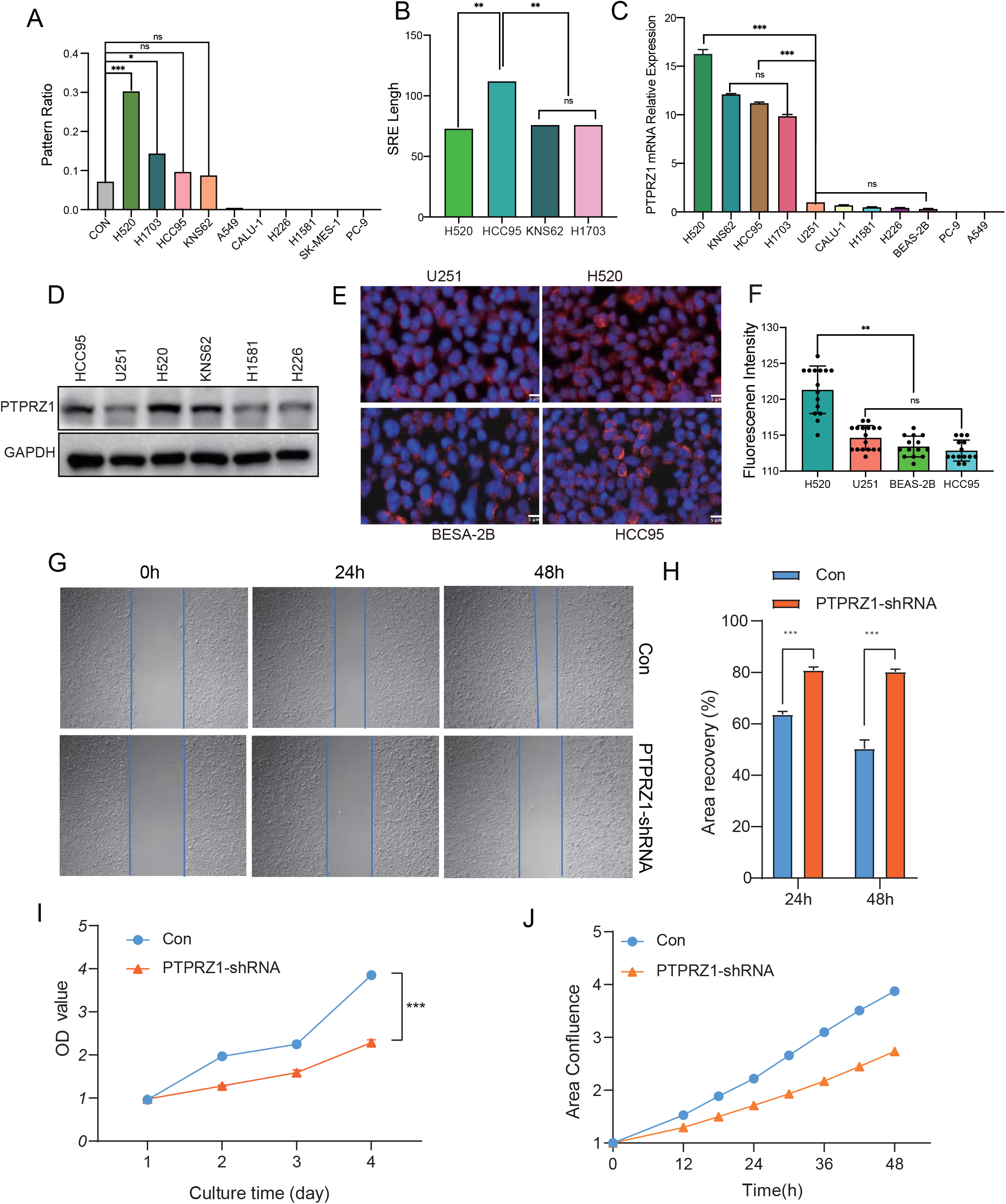
Highly expressed *PTPRZ1* could promote the proliferation and migration of tumor cells in vitro. A. Detection of the SRE pattern ratio by targeted sequencing in ten NSCLC cell lines (pattern ratio = count of reads harboring the SRE/count of reads mapped to the target region chr7: 121603000-121604000, the pattern ratio of 0.06% served as a control, ≥ 0.06% indicates the presence of the SRE). B. The length of the SRE predicted by the local assembly of targeted sequencing data from H520, HCC95, KNS62 and H1703 LUSC cells. C. The relative expression levels of *PTPRZ1* determined using RT‒PCR in 11 cell lines. D. Western blot analysis of *PTPRZ1* expression in HCC95, U251, H520, KNS62, H1581,, and H226 cells. E. Fluorescent immunostaining for the *PTPRZ1* protein in the LUSC cell line H520, HCC95, glioma cell line U251 and normal bronchial epithelial cell line BEAS-2B cells. Staining in panels is representative of three samples. Nuclei (DAPI) are stained blue. Scale bars: 5μm. F. Fluorescence intensity statistics of *PTPRZ1* protein expression levels in H520, HCC95, U251 and BEAS-2B cells are shown in Figure. 4E. Unpaired two-tailed t tests were used to compare two groups. Error bars represent the mean ± standard error of the mean. G. Wound healing detected cell migration ability after PTPRZ1 knockdown by shRNA in HCC95 cells. H. The relative migration ability of HCC95 cells and PTPRZ1 knockdown HCC95 cells shown in Figure. 4G (n = 3). ***P < 0.001. Student’s t test was used. I. CCK-8 detected the proliferation rate of cells in HCC95 cells and PTPRZ1 knockdown HCC95 cells at 1-4 day (n = 3). J. The confluency of HCC95 cells and PTPRZ1 knockdown HCC95 cells was monitored for 48h by incuCyte system to detect cell proliferation rate.

To further investigate the potential function of PTPRZ1 in LUSC cells, we knocked down the PTPRZ1 gene in HCC95 cells by shRNA. Bifunctional investigations revealed that the knockdown of PTPRZ1 resulted in suppressed LUSC cell migration and proliferation activity (Figure 4G-4J). As shown, the cells migration ability of the PTPRZ1-shRNA group was suppressed compared with the Control group (Figure 4G-4H). The viability and proliferative ability of the HCC95 cells was significantly inhibited after PTPRZ1 knockdown (Figure 4I-4J). These results suggested PTPRZ1 promote cell migration, proliferation, and the carcinogenesis of LUSC cells. Next, we generated lentiviral constructs in PTPRZ1-overexpressing H226 cells (Figure S4J-S4N), and we could find the expression of c-MYC, Cyclin-D1 and E-cadherin increased following PTPRZ1 overexpressing. The results showed that the PTPRZ1-OE group substantially increased LUSC cell migration and invasion ability (Figure S4M-S4N). Taken together, these results support the notion that this SRE is a tumor-activator element in LUSC and that certain somatic mutations could promote PTPRZ1 activity to regulate the proliferation, migration and invasion of tumor cells.

### Somatic Transposable Elements in NSCLC Genomes Are Mainly Composed of Complex Structures

In total, 4574 somatic TEs were identified in our NSCLC cohort. The identified somatic TEs are of high fidelity as we only applied reads that completely span these TEs. To further determine the sequence of insertion sequence, we performed polishing using insertion-supporting reads to achieve a corrected consensus sequence for TEs and their flanking sequences (Methods, Table S4). The average length of the assembled contigs was 25 kb (645 bp to 127 kb, median 24 kb, Figure S5A). Through the annotation of somatic TEs (Methods), we found that 55% of the somatic TEs had complex structures (Figure 5A). These somatic TEs consisted of a complex combination of multiple TE segments, which we named “complex transposable elements” (complex TEs). An example of a complex TE is shown in Figure S5B, representing somatic complex TE with a length of 1155 bp is composed of SINE/AluY, LINE/L1MEi, SINE/AluSx and unmasked sequences. Somatic complex TEs were longer than somatic solo TEs on average (the average length of complex TEs was 1037 bp, whereas the average length of solo TEs was 311 bp (Figure S5C). Similar to what has been reported for nonallelic homologous recombination (NAHR) in previous studies, some complex TEs were composed of 2 SINEs (10.34%) or 2 *LINEs* (6.50%). However, in addition to these NAHRs, our analysis revealed that the majority (83.16%) of complex TEs also contained other repeat sequences (Figure 5B). Although the aberrant structures varied, 90.38% of somatic complex TEs contained SINEs or LINEs, indicating that SINEs and LINEs dominated the landscape of somatic complex TEs in the NSCLC genome. Moreover, we found that complex TEs were predominantly composed of the youngest TE subfamilies, such as AluY, L1HS, SVA (SINE-VNTR-Alu), and HERV-K (human endogenous retrovirus type K). These findings suggest that these still-active TE subfamilies in the human genome may be involved in complex TE formation (Figure S5D).

**Figure 5.**
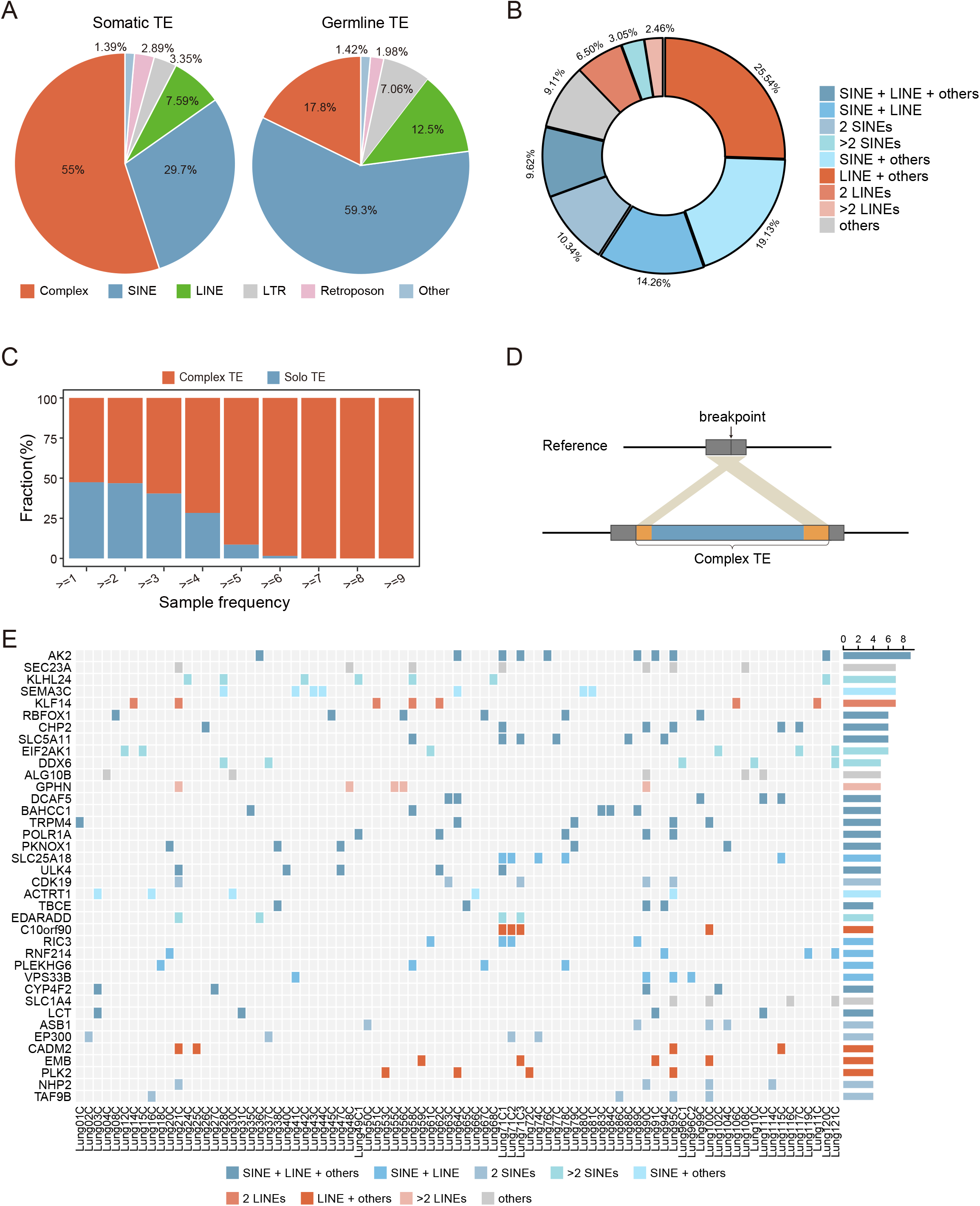
Somatic complex TEs may frequently affect the NSCLC genome. A. Pie charts depict the proportions of TE types in somatic TEs (left) and germline TEs (right). B. The composition of the complex TEs. Repeats other than SINEs and LINEs are classified as ‘others’. C. The proportions of somatic solo TEs (blue) and somatic complex TEs (orange) in various numbers of shared samples are shown. The fraction of complex TEs increases as the shared number of samples increases. D. A schematic representation of homologous sequences at both ends of somatic complex TEs. Black lines, reference sequence; gray blocks, homologous sequences on reference; red and blue blocks, complex TE. E. New open chromatin regions related to recurrent complex TEs and their closest genes. The oncoprint shows the contribution of complex TEs across samples. Color indicates the different TE composition types. The bar chart (right) shows the shared number of samples.

Complex TEs accounted for 55% of all somatic TEs, although complex TEs represented only 17.80% of germline TEs (Figure 5A). In all patients, complex TEs represented a higher proportion of somatic TEs than germline TEs (Fisher’s exact test, one-sided, q value < 0.1, Figure S5E). Moreover, complex TEs were the major type of somatic TE shared among samples, and the proportion of somatic complex TEs increased as the number of shared samples increased (Figure 5C). Therefore, it is likely that somatic complex TEs play functional roles in the NSCLC genome.

To identify the potential mechanism responsible for the formation of somatic complex TEs, we analyzed the flanking sequences around breakpoints. We observed that 68.17% (1298/1904) of the somatic complex TEs were flanked by a TE on at least one side. In previous studies, homologous repeats around the breakpoint have been used to infer potential mechanisms of origin of structural variations (Fujimoto et al., 2021). To determine whether breakpoint junctions contain homologous sequences, somatic complex TEs were aligned to corresponding upstream and downstream flanking assembled sequences separately. Interestingly, we observed that 92.84% (1205/1298) of these complex TEs followed a pattern in which the somatic inserted TE segments contained flanking homologous sequences at one end at least (Figure 5D). The presence of flanking homologous sequences indicates that somatic complex TE formation could potentially be mediated by recombinational repair of TEs. The integration of multiple additional fragments may be triggered by a strand invasion mechanism during the repair process, similar to previous observations of certain chimeric LINE insertions (Gilbert et al., 2002). Furthermore, the vast majority of the complex TEs shared by different samples (83.22%, 362/435) were flanked by TEs, among which 97.24% (352/362) presented a homologous sequence pattern. We identified a significant enrichment of SINEs in the homologous sequences of these shared complex TEs (Figure S5F, chi-square test, p value = 5.9e-06), suggesting that the occurrence of complex TEs shared by different samples was likely related to SINE-mediated recombination.

A total of 99.54% of the somatic complex TEs shared by different samples were inserted in noncoding regions. These complex TEs may impact the gene regulatory network by affecting chromatin accessibility. We identified somatic complex TEs that were related to new open chromatin regions by jointly analyzing genomic and ATAC-seq data (Methods). In total, 34.48% (150 out of 435) of the complex TEs shared by different samples were related to new open chromatin regions at the sites of their insertion, among which 38 events were shared by at least 4 samples. Compared with all somatic complex TEs, the complex TEs shared by different samples harboring SINE segments were more likely to be associated with new open chromatin regions (Figure S5G, Fisher’s exact test, one-sided, p value < 0.05). Furthermore, we identified 38 genes adjacent to the complex TEs shared by different samples (≥ 4 samples shared) that were related to new open chromatin regions (Figure 5E, S5H). These genes included key cancer-associated genes, such as *DDX6*, *GPHN*, *EP300*, *EIF2AK1*, and *SEMA3C* [52–57].

### Somatic TEs Affect the NSCLC Transcriptome Through Expression

Previous studies have reported that TE copies can function not only as regulatory elements but also as a source of alternative splicing in gene expression [58]. The expression of a TE itself is one of the alternative splicing mechanisms caused by TEs [24, 59] and is a process that can provide raw material for the emergence of new isoforms, which may take on important cellular functions in tumors [60]. However, this phenomenon of somatic TEs has been understudied in cancer research. According to our data, more than 68% of somatic TEs were located in the gene body or the 5 kb flanking regions of the gene, which were likely to be expressed or expressed within flanking genes. To further explore this hypothesis, we mapped the RNA-seq reads of each patient to the corresponding assembled contigs (Methods). A total of 31.78% (1228/3864) of the somatic TEs were expressed in tumor samples. Specifically, somatic TEs were determined to be expressed under two conditions: at least one RNA-seq read could align to and span the breakpoints, or split RNA-seq reads could be mapped to both the somatic TEs and their adjacent genes. We validated somatic TE expression events in one available tumor sample using RT‒PCR (Figure S6A, Methods), and these TE expression events also changed the protein sizes encoded by the inserted genes (Figure S6B). A vast majority (76.19%) of the tumor samples showed at least one expression event (ranging from 1 to 262, median 3), indicating that the expression of somatic TEs is widespread in NSCLC. RNA-seq data for the 1228 TEs demonstrated the presence of somatic expressed TEs in 1128 genes (Figure 6A), among which 11.44% (129 out of 1128) were annotated by the Catalog Of Somatic Mutations In Cancer (COSMIC) or The Network of Cancer Genes (NCG) as cancer-related genes, whereas 28 genes were oncogenes (Figure 6A, Table S5A). A total of 167 genes harbored expressed TEs in at least 2 patients. The most frequent expression was identified in a cancer-related gene, *LRP1B* (in 6 patients), encoding a putative tumor suppressor in NSCLC, gastric cancer and neuroendocrine carcinoma [61, 62]. The expressed TE-related genes were enriched in cancer-related pathways, such as the Rap1 signaling pathway (q value = 0.011), ErbB signaling pathway (q value = 0.022), and chemokine signaling pathway (q value = 0.027) (Figure 6B, Table S5B). Cancer-related Gene Ontology (GO) terms were also significantly enriched (q value < 0.05), such as cell division (q value = 0.00087, 22 genes including 2 COSMIC genes, *ZFYVE19* and *KNSTRN*) and cell migration (q value = 0.0015, 17 genes including 3 known lung cancer-related genes, *ERBB4*, *EPHA3*, and *ARHGAP35*) (Table S5C).

**Figure 6.**
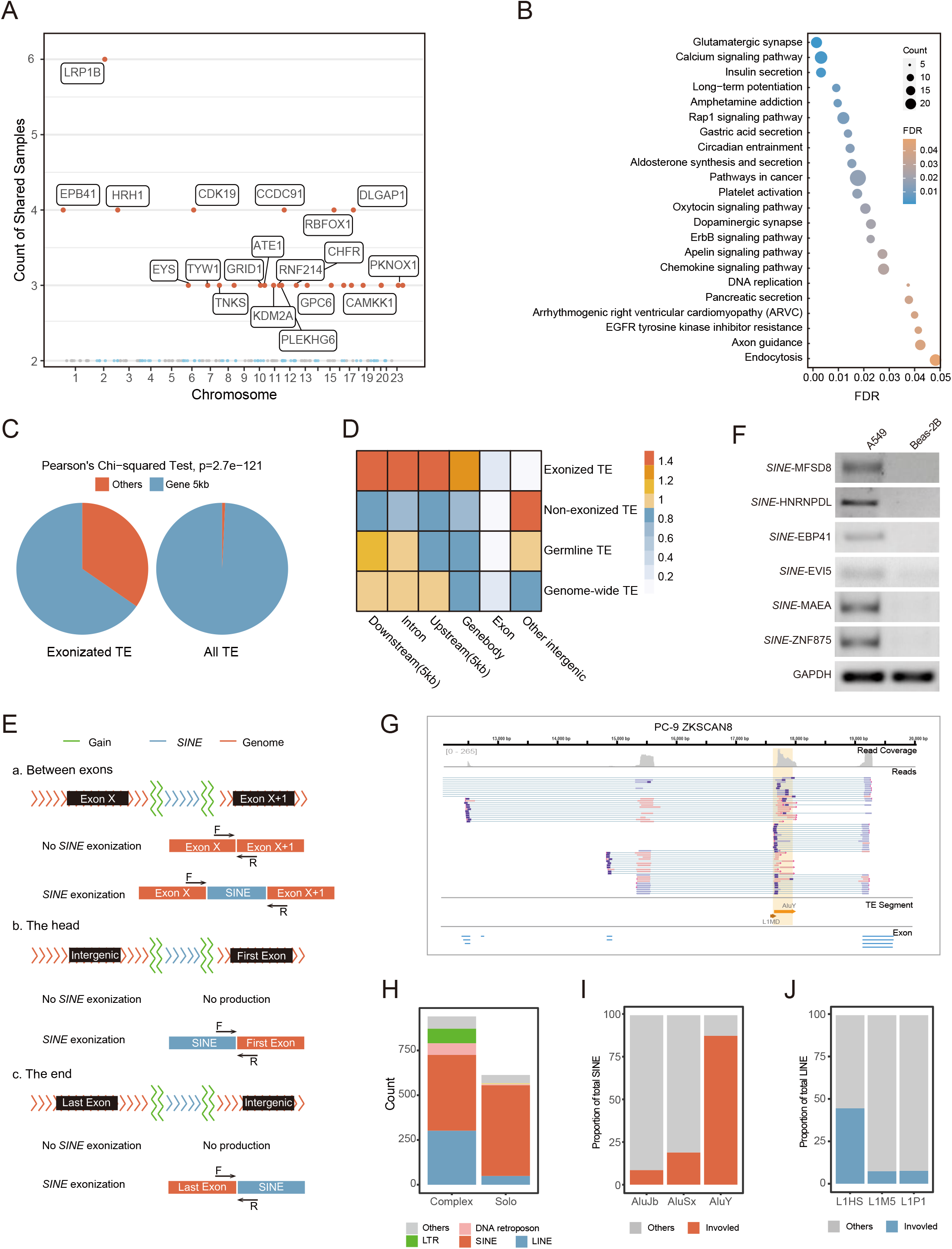
Detection and validation of somatic expressed TEs. A. Manhattan plot for genes harboring somatic expressed TEs. Chromosomes are represented on the x-axis. The shared number of samples is represented on the y-axis. Orange dots represent genes shared by at least 3 tumor samples. The names of protein-coding genes are annotated near the corresponding dot. B. Enrichment map showing *KEGG* pathways in which expressed TE-related genes were enriched. C. Genome-wide insertion site composition of somatic expressed TEs (‘expressed TE’) and all somatic TEs (‘all TE’). These TEs were grouped as ‘Gene 5 kb’: inserted within the gene or the 5 kb region adjacent to the gene, and ‘Others’: inserted within intergenic regions excluding 5 kb regions adjacent gene. D. Enrichment of somatic expressed TEs in genomic elements. Color indicates the odds ratio range, as in the legend. No clustering is presented. E. Schematic illustrating the location of the Alu insertion and the location of the primers used to identify Alu externalization events. There are three types of Alu insertion events: between exons (a), in the head (b) or at the end of the genes (c). The blue line indicates the Alu, and the green line indicates where the Alu is inserted. F represents the forward primer, and R represents the reverse primer. The lengths of the PCR products were all less than 300 bp, excluding the length of the TE. F. Agarose gel analysis of SINE expression in the A549 cell line. The Beas2B cell line was used as the control. SINE, short interspersed nuclear element. G. Genome Browser views displaying the primary alignments overlapping the sequences of PCR-validated expressed TEs and their adjacent exons of *ZKSCAN8*. The light-yellow-shaded area shows the insertion, while line segments present the segments constituting the insertion, different colors represent different types of TE families, and arrows represent the direction of each TE segment. H. Counts of different TE families involved in somatic expressed TEs. Somatic expressed TEs were grouped into complex TEs and solo TEs. I and J. Proportion of somatic expressed TEs involving Alu and L1. The top three subfamilies of Alu or L1 that had the highest counts are shown.

Compared with all somatic TEs, somatic expressed TEs were more frequently located in the gene body and regions adjacent to the gene body (1219/1228, 99.27%, Figure 6C), especially in promoters and introns (hypergeometric test, Benjamini‒Hochberg-corrected q < 0.01, Figure 6D). A total of 74.23% of somatic expressed TEs were located in gene introns, providing new alternative splicing sites and offering potential for intron retention. A total of 11.31% of somatic expressed TEs were located upstream of genes (ranging from 75 bps to 4976 bps, median 2093 bps) and may act as potential alternative promoters of these genes in NSCLC (Table S5D). A total of 10.19% of somatic expressed TEs were located downstream of genes (ranging from 2 bps to 4950 bps, median 1901 bps), providing potential alternative transcription end sites. Only 4.27% of somatic expressed TEs were located in other regions.

To further verify the somatic TE expression events, we analyzed long-read transcriptome (LRS RNA-seq) data from 3 lung carcinoma cell lines (A549, H1299 and PC-9) [11, 63]. A total of 288 TE expression events were included in the RNA-seq data (111, 111 and 66 from A549, H1299 and PC-9 cells, respectively, Table S5E, Methods). These TE-expressed transcripts were supported by at least 3 LRS RNA-seq reads spanning both a TE and one or more exons of associated genes. Twenty percent of TE expression events were shared by at least 2 cell lines, showing that they were widespread in NSCLC tumor cells (Figure S6C). Furthermore, retrotransposition-PCR (RT‒PCR, Figure 6E) was used to assess 21 SINE expression events (6, 10 and 8 from A549, H1299 and PC-9, respectively; Figure 6F and S6D). We sequenced all the PCR-validated TEs of the PC-9 and H1299 cell lines using the QNome-3841 platform and confirmed that all TEs matched their expected expressed TEs identified from our dataset (Table S5F). All somatic TEs were supported by at least 15 LRS RNA-seq reads, which mapped to both somatic TEs and the exons of associated genes. These results confirmed the presence of our predicted TE-expressed events, although some TEs exhibited low expression. Moreover, SINEs were the predominant type of expressed TE segments (Figure 6G, S6K). The sequence of the full-length SINE inserted into *ZKSCAN8* was obtained by targeted sequencing, and the SINE insertion changed the size of the protein encoded by *ZKSCAN8* in PC-9 cells (Figure S6E). To investigate the underlying biological role of SINE expression in gene expression and cell proliferation in cell lines, we overexpressed ZKSCAN8^WT^ and ZKSCAN8_SINE_ in the Beas-2B cell line using a recombinant eukaryotic expression plasmid. First, we confirmed overexpression of ZKSCAN8_WT_ and ZKSCAN8_SINE_ in Beas-2B cells compared with the control cell line infected with empty vector (Beas-2B-Vector), the presence of the plasmids containing the SINE sequence led to increased transcription levels of the *ZKSCAN8* gene (Figure S6F and S6G). Then, we detected ZKSCAN8 protein expression by Western Bolt and observed that Beas-2B cells transfected OE-ZKSCAN8_SINE_ plasmids expressed an additional band larger than 65 kDa (Figure S6H), similar to the results in PC-9 cell (Figure S6E). We further explored the role of SINEs in carcinogenesis. We observed greater proliferation rate of Beas-2B-OE-ZKSCAN8_SINE_ cells than Beas-2B-OE-ZKSCAN8_WT_ cells or the control cell lines according to the Cell Counting Kit (CCK8) proliferation assay (Figure S6I). Accordingly, the migration capacities of Beas-2B-OE-ZKSCAN8_WT_ and Beas-2B-OE-ZKSCAN8_SINE_ cells were evaluated via wound-healing assays. We found that the wound closure rate of Beas-2B-OE-ZKSCAN8_SINE_ cells was significantly higher than that of Beas-2B-OE-ZKSCAN8_WT_ cells and the corresponding control cell lines (Figure S6J). Taken together, these findings implied that ZKSCAN8-expressed SINEs can significantly upregulate gene expression, change protein size, and stimulate cell proliferation, thereby presenting the potential to facilitate tumor progression.

Previous works have demonstrated that long terminal repeats (LTRs), DNA transposons and LINEs are commonly overexpressed TE subfamilies in the cancer genome. Indeed, we identified 369 somatic expressed LINE segments, 88 somatic expressed LTR segments, and 70 somatic expressed DNA transposons (Figure 6H). Several studies [27, 64] have reported that LINEs dominate somatic retrotransposition in multiple cancer types, whereas SINEs are a minor category of somatic expressed TEs. Surprisingly, our results showed that 75.73% (930/1228) of somatic expressed TEs contained SINE segments, among which the number of solo SINEs was 1.2-fold that of complex TEs. Such a large proportion of somatic expressed SINEs was not reported previously. In contrast, the majority of somatic expressed LINEs (320/369, 86.72%), LTRs (82/88, 93.18%), and DNA (65/70, 92.86%) were involved in complex TEs. In summary, our results demonstrated that SINEs overwhelmingly dominate the landscape of somatic expressed TEs in the NSCLC genome.

Consistent with previous findings [65], we found that younger TEs were more likely to be expressed in the NSCLC genome. The number of *AluY* (87.85%) expression events was approximately 4.6-fold higher than that of the older subfamily *AluSx*, which ranked second in the number of expression events (Figure 6I). *AluY* is the youngest subfamily of *Alu*, which exhibits the greatest ability to move in the human genome [66]. *L1HS*, the youngest LINE subfamily, which is human specific, comprised 44.72% of expressed LINEs (Figure 6J). The number of *L1HS* elements was approximately 5.8-fold higher than those of the older subfamilies *L1M5* and *L1P1*. These findings confirmed the previous finding that the TEs that are overexpressed in cancer are the evolutionarily youngest TEs [65].

## Discussion

In this study, we demonstrated the landscape of high-confidence somatic INSs (at least 50 bp in length) in NSCLC genomes by generating a substantially large long-read sequencing (LRS) dataset (17.29 Tb) from 242 patients. The length of LRS reads allows the determination of the complete sequence and the flanking region of INSs, even in highly repetitive and variable regions. The relatively high sequencing error is a major challenge for downstream analysis. For example, the popular aligners and SV callers for LRS based on the seed-chain-align strategy [31, 67] lead to many problems when calling INSs, given the irregularity of breakpoints on the supporting reads of one INS and the difficulty of detecting repeat INSs. To address these issues, we developed a statistical model to fine tune the breakpoints of INSs and manually examined all candidate somatic INSs. During the manual examination, we filtered out candidate somatic INSs that were nested with other SV types, and INSs exhibited variable lengths within the same loci. Our study represents a step in the development of a novel computational approach that detects somatic SVs based on an LRS dataset. One caveat is that FFPE tissue is not recommended for LRS, to ensure the integrity and length of DNA and avoid the artifacts caused by the deamination of cytosine bases to uracil in FFPE tissues. The use of fresh or frozen samples is recommended whenever possible.

One of the advantages of LRS was that it enables the reliable identification of TDs, which is impossible using NGS short reads. We identified an LUSC-enriched somatic SRE between the distal enhancer regions targeting the *PTPRZ1* gene. Although the SRE could also be identified in SRS data by ExpansionHunter Denovo, the number of supporting reads was low (below 3) when 50% of the samples contained the SRE (Figure S2C). In contrast, while the sequencing depth of LRS data was lower than that of SRS data (28.5X versus 34.67X), many more supporting reads were identified in LRS data. This advantage enabled us to set a strict threshold for screening somatic INSs. During the analysis of LRS data, we required a minimum number of 3 supporting reads, which is why the SRE was identifiable in two samples based on SRS data but not LRS data.

We experimentally validated the interaction between the SRE and *PTPRZ1* using both the Hi-C and 3C-PCR methods. Moreover, we showed that the SRE promoted *PTPRZ1* expression, and the rescue experiment of *PTPRZ1* in LUSC H226 cells, which do not contain the somatic SRE, indicated that *PTPRZ1* could promote cell proliferation and migration. These results suggested that *PTPRZ1* might be a new target for the diagnosis and treatment of LUSC. However, more experiments are needed to elucidate the mechanisms and therapeutic value of *PTPRZ1* in LUSC. LUSC patients receive limited benefits from targeted diagnosis and treatments due to the lack of LUSC-enriched genomic markers and actionable gene targets [4, 68]. *PTPRZ1* is a potential bridge to fill this gap.

With the LRS data, we were able to examine the full sequence of the somatic TEs in the lung carcinoma samples. We demonstrated that more than half of the somatic TEs were complex TEs that consisted of multiple TE segments. Compared with germline TEs, somatic TEs had a higher proportion of complex TEs, indicating that TEs are more active in tumor cells. We also characterized the genomic context of complex TEs. In total, 68.17% of the complex TEs were inserted into germline TE loci through a possible recombination mechanism. Using ATAC-seq data, we also showed that complex TEs shared by different samples affected gene regulation by changing the chromatin state. Larger studies including more patients harboring complex TEs shared by different samples and in vitro or animal models are needed to further verify this hypothesis.

Another advantage of LRS data is that it enabled the accurate identification of the insertion locations of somatic INSs. Our study revealed extensive somatic TE expression events in NSCLC that occur in pivotal cancer-related genes, such as *ERBB4* and *EBP41*. The expression of TEs, particularly those of the LINE-1 and LTR subtypes, has been previously explored [11, 64, 65, 69, 70]. However, few studies have reported somatic SINE expression as a major mechanism of abnormal transcriptome changes in NSCLC, likely because previous studies were limited by the use of SRS data. Due to the high repetition of TEs in the human genome, SRS reads originating from TEs can often be mapped equally well at various positions in the genome. Although SINEs are approximately 300 bp long, the number of SINEs in the human genome is over four times that of LINEs. Additionally, Younger TEs are nearly identical to each other. Even with 2 x 100 bp paired-end libraries, less than half of the reads emanating from the youngest TEs are informative [71]. As a result, the measurement of young SINE expression and the understanding of how they regulate gene expression have been limited by the use of SRS technology. Increasing the read length can help mitigate these effects, as longer reads provide more information about the flanking sequences, thereby aiding in the accurate detection of SINEs. In fact, recent large-cohort studies using LRS data have revealed an unprecedented number of germline SINE insertions [13, 14]. The LRS data we generated herein provided the opportunity to reveal previously hidden somatic expressed SINEs in NSCLC genomes. Indeed, we found that SINEs, especially those of the Alu subtype of the young lineage, dominated somatic expressed TE events. The mechanisms underlying this predilection about TE expression in certain malignancies are poorly understood. Overall, the types of tumors that acquire somatic TE insertions correspond to those types that show TE protein expression (long interspersed element-1 protein expression is a hallmark of many human cancers), and controls on expression are probably important. Here, we reported that SINE expression could alter the gene expression levels (Figure S6G) and the sizes of the proteins encoded by genes harboring insertions (Figure S6F, S6H). It is likely that the SINE expression events we reported here are still an underestimation because we used short-read RNA-seq data in the analysis. A comprehensive long-read RNA-seq dataset would certainly provide a more accurate picture.

*PTPRZ1* encodes a transmembrane protein tyrosine phosphatase that is highly expressed in SRE-enriched LUSC. Gene fusions were found to be the most common type of mutation [72]. *PTPRZ1-MET* fusion can be identified in 8%-16% of glioma patients and is associated with a poor prognosis (Bao et al., 2014). *PTPRZ1-MET* fusion is a predictive biomarker for treatment with MET kinase inhibitors (Matjasic et al., 2020). Given the strong upregulation of *PTPRZ1*, the antagonization of *PTPRZ1* expression and/or signaling may be a promising strategy for inhibiting tumor growth. Several inhibitors of *PTPRZ1*, including MY10, MY33-3 [73] and NAZ2329 [74], have been identified. For example, NAZ2329, a cell-permeable small molecule that allosterically inhibits *PTPRZ1*, reduces the expression of *SOX2* and abrogates the sphere-forming abilities of glioblastoma cells. However, neither *PTPRZ1* nor corresponding antibodies have been translated into clinical practice, and the most effective drugs for use in this context are not clear. Therefore, the possible role of *PTPRZ1* in diagnosing and treating LUSC should be considered in the future.

In summary, by generating a large genomic dataset with emerging LRS technology and discovering many previously unidentified somatic INSs in the NSCLC genome, this study represents an important effort to fill existing knowledge gaps and provides an opportunity to comprehensively explore the functional impacts of a new type of somatic alteration in NSCLC.

**Figure S1.**
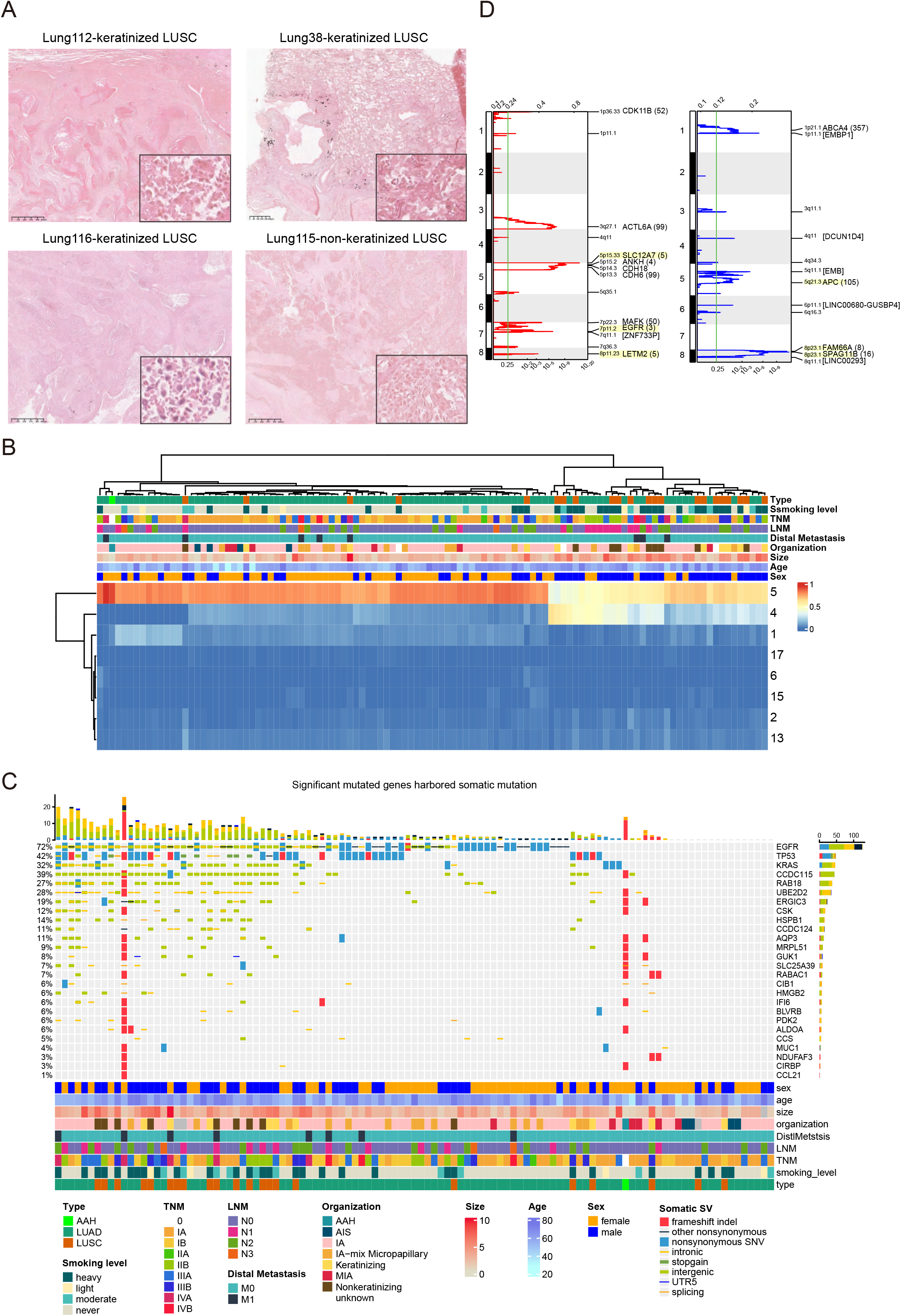
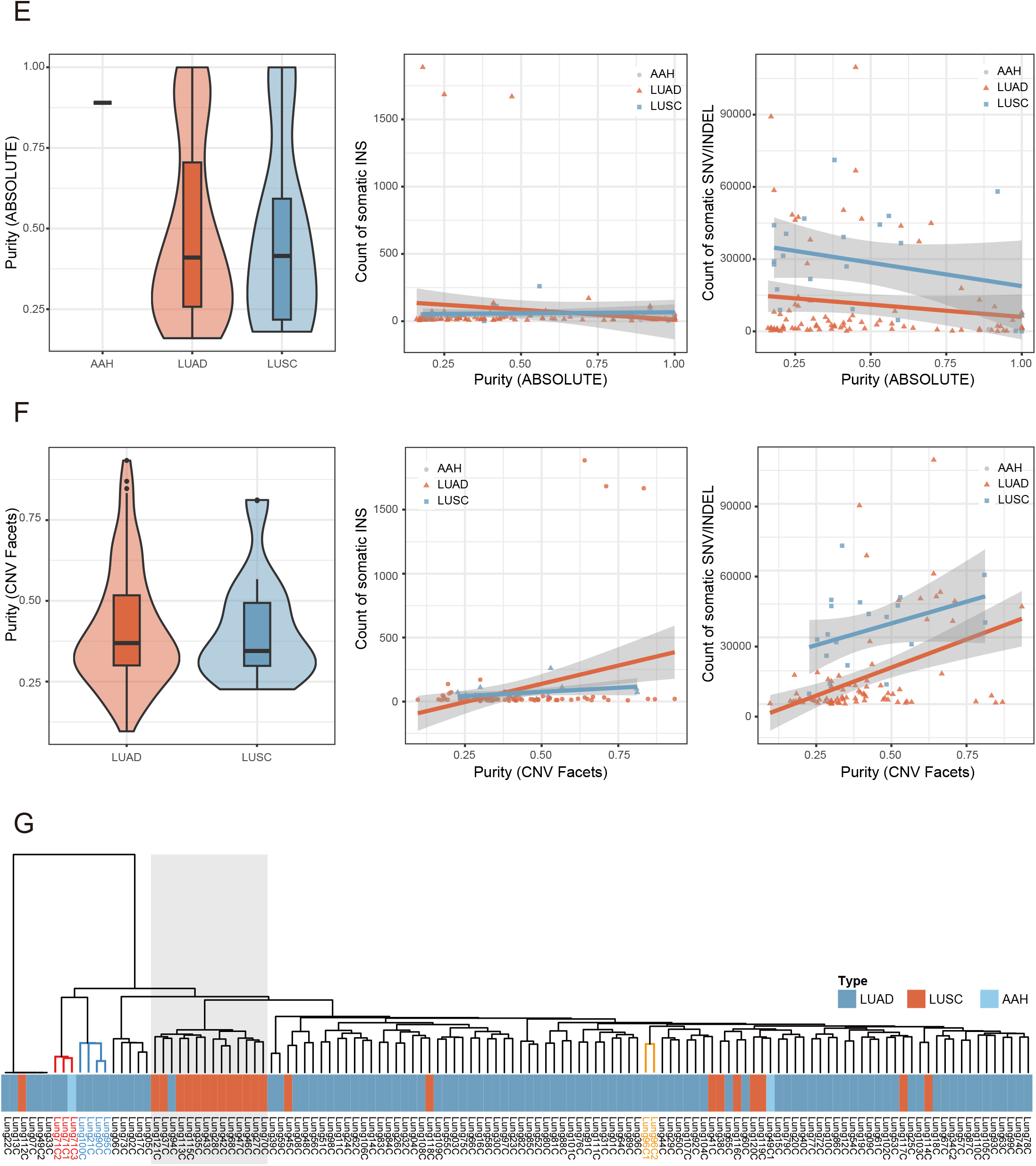
Characteristics of somatic INSs and SNVs/indels in NSCLC. A. LUSC classified as keratinized and nonkeratinized squamous were annotated by pathologists. B. Heatmap showing the hierarchical clustering of NSCLC samples based on the somatic mutation signature, with the clinical information of the samples. C. Co-mutation plot obtained from the short-read whole-genome sequencing of 110 NSCLC cases. Significant genes and driver genes of LUAD and LUSC, which were reported by TCGA studies, are shown and ranked in order of decreasing prevalence. D. Somatic focal amplification regions and deletion regions identified by GISTIC with the q value at bottom. Significant regions (q value < 0.05) are annotated with one of the genes contained in the peak, with the number of genes in the peak in parentheses. Regions previously reported in TCGA are highlighted in yellow. E. The left panel shows no significant differences in ABSOLUTE purity between LUAD and LUSC (p>0.05, Wilcoxon rank-sum test). In the middle panel, there is no significant correlation between ABSOLUTE purity and the count of INSs (r = 0.0236, p > 0.05, Spearman correlation). Likewise, the right panel demonstrates no significant correlation between ABSOLUTE purity and the SNV/indel count (r = −0.125, p > 0.05, Spearman correlation). F. The left panel shows no significant differences in CNV Facets purity between LUAD and LUSC (p>0.05, Wilcoxon rank-sum test). In the middle panel, there is no significant correlation between CNV Facets purity and the count of INSs (r = 0.0497, p > 0.05, Spearman correlation). Likewise, the right panel demonstrates a significant correlation between ABSOLUTE purity and the SNV/indel count (r = 0.3824, p = 4.806e-05, Spearman correlation). G. Heatmap showing the hierarchical clustering of NSCLC samples based on somatic insertions is shown with the clinical information of the samples (the gray area indicates a cluster mainly consisting of LUSC samples, the sample IDs highlighted in red were obtained from patient Lung71, the sample IDs highlighted in orange were obtained from patient Lung96, and the sample IDs highlighted in blue were classified in a subclass with a high SV burden).

**Figure S2.**
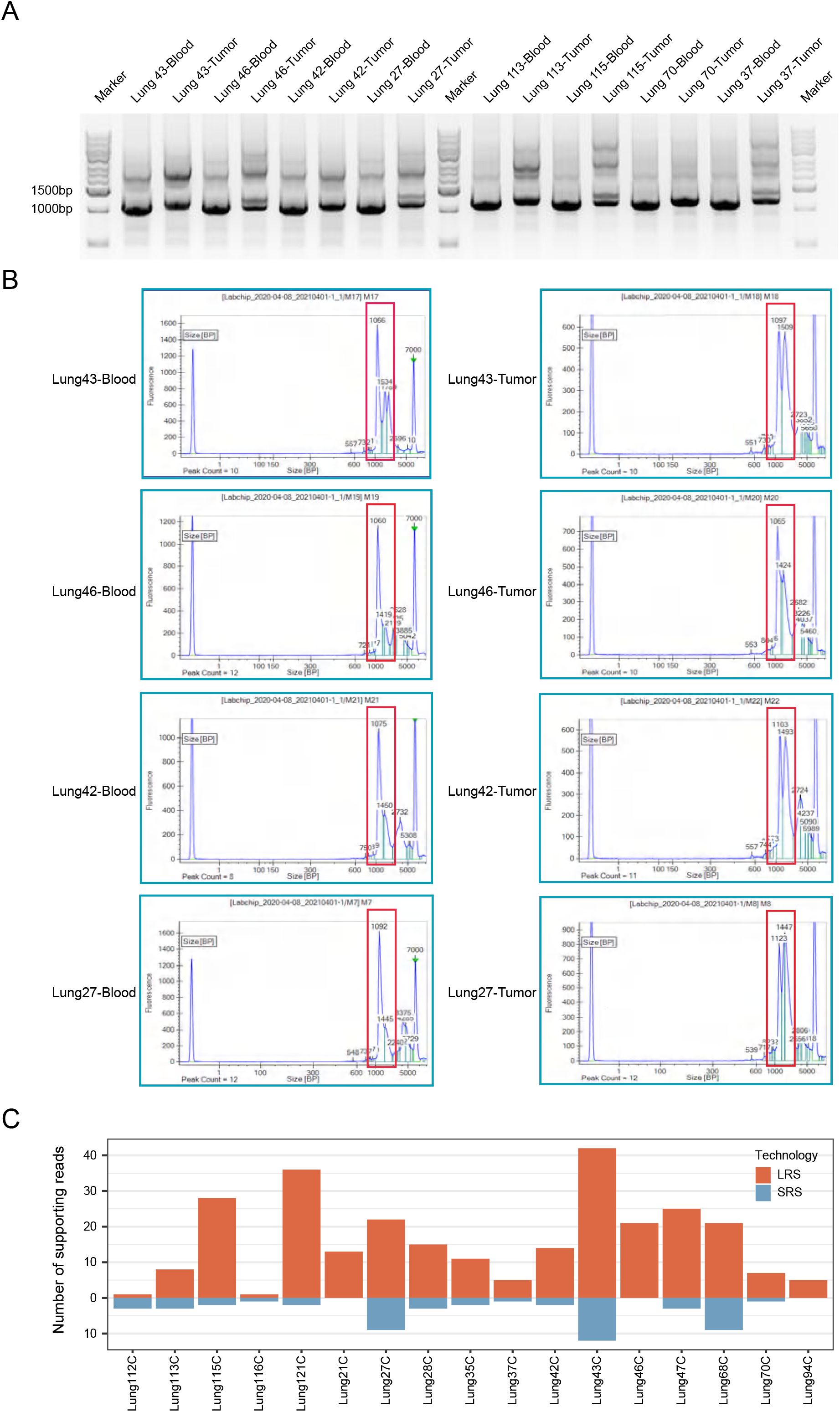
Identifying SREs in blood and tumor tissues of LUSC patients by gel electrophoresis, capillary electrophoresis and SRS. A. Horizontal gel electrophoresis of SRE-harboring DNA fragments from blood and tumor tissues, obtained by the targeted sequencing of LUSC patients. B. Size distribution of SRE-harboring DNA fragments in blood and tumor tissues evaluated by LabChip GX capillary electrophoresis in LUSC patients. C. Number of reads supporting the SRE identified based on SRS and LRS data. The SRE was not considered to be identified in Lung112C or Lung116C based on LRS data, as the number of supporting reads was below 3.

**Figure S3.**
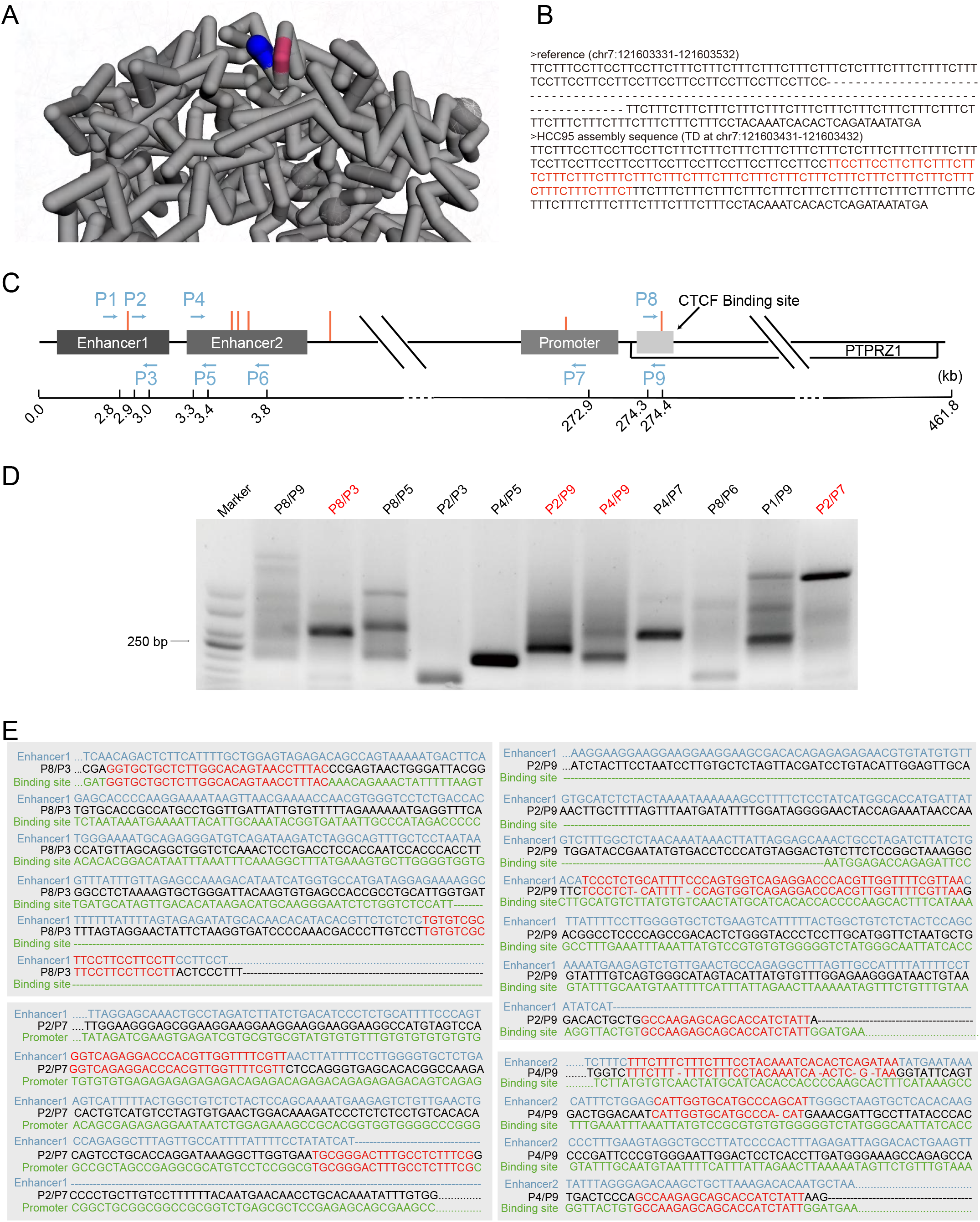
The LUSC-enriched SRE may contact distal enhancers and *PTPRZ1* and exhibit high expression in SCC. A. The predicted 4D nucleosome structure around the LUSC-enriched SRE and *PTPRZ1*. The region that harbors the SRE and distal enhancers is highlighted in blue, and the region that harbors *PTPRZ1* is highlighted in pink. B. The reference sequence of the region where the somatic SRE was inserted (top) and the assembled SRE sequence based on the long-read targeting region sequencing data of HCC95 cells are shown (bottom). Red letters represent the SRE sequence of HCC95 cells. C. Paired primers (P1–P9) containing two control test primer pairs (P2/the P3, P4/P5) were designed for the detection of cis-acting interactions. Small vertical bars in red represent MseI digestion sites. Blue arrows show the primers. D. The PCR products were detected by agarose gel electrophoresis. Red marks indicate that the sequencing results including expected chromatin interactions. E. PCR products of these primer pairs were sequenced, and 4 products matched the expected chromatin interaction. Matched sequences are labeled in red.

**Figure S4.**
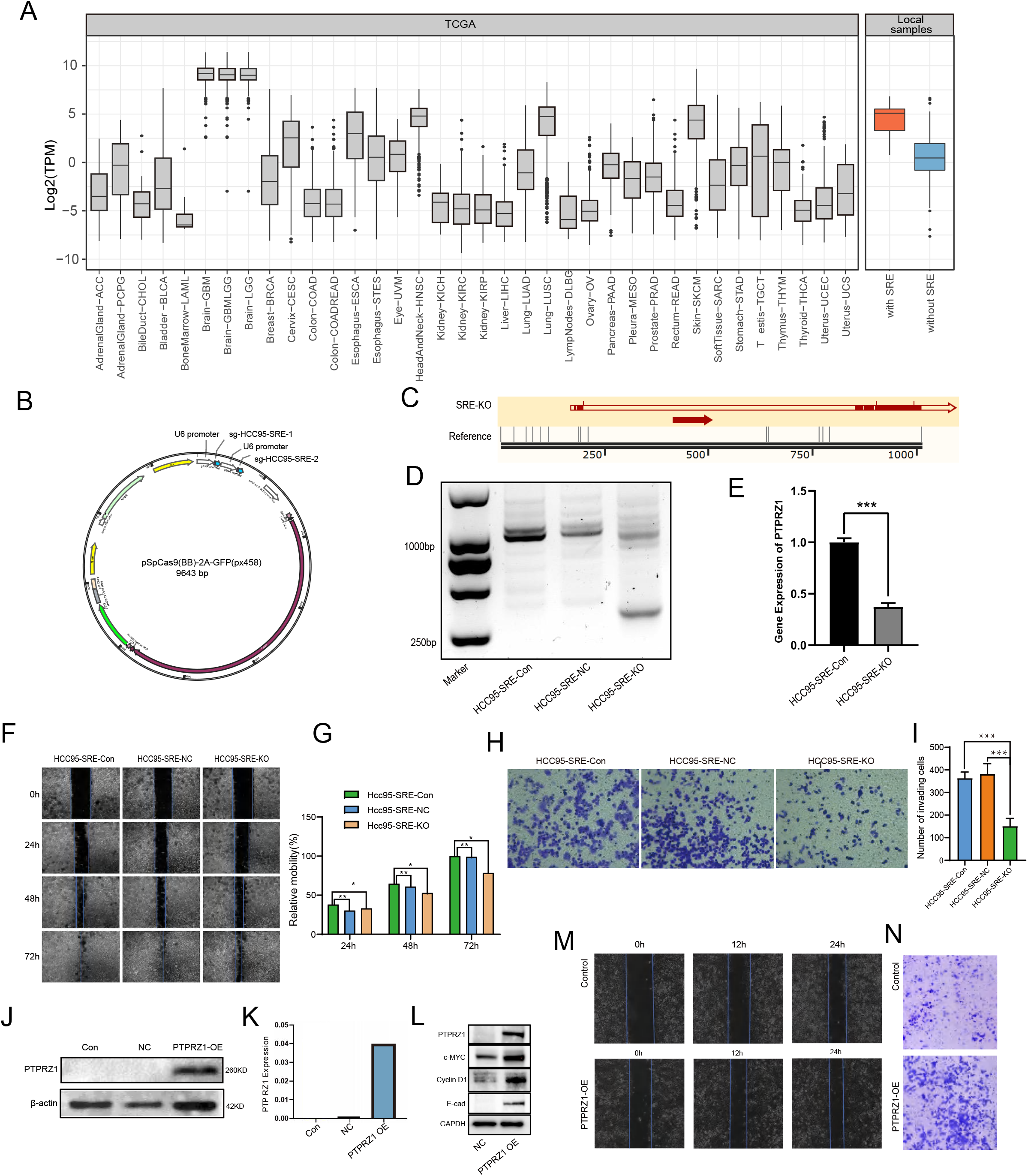
SRE regulates *PTPRZ1* gene expression in LUSC cell lines, and overexpression of *PTPRZ1* increases LUSC cell migration and invasion ability. A. Boxplot showing *PTPRZ1* expression in TCGA cancer samples and NSCLC samples with LUSC-enriched SRE. B. Profile of HCC95 SRE-KO plasmid. C. Sequence alignment of the HCC95 SRE-KO cells compared with the HCC95 genomic reference sequence (SRE was indicated by a red arrow). D. Agarose gel electrophoresis of PCR amplicons products (HCC95 SRE-Con, HCC95 SRE-NC, HCC95 SRE-KO). E. The relative expression levels of *PTPRZ1* determined using RT‒PCR in HCC95 cells and HCC95-SRE-KO cells F. The cell migration ability of HCC95 cells and HCC95-SRE-KO cells was measured in cell scratch assays at 24 h, 48 h, and 72 h. G. Graph of the relative cell migration ability of HCC95 cells and HCC95-SRE-KO cells is shown in Figure. S4F. H. The cell invasion ability of HCC95 cells and HCC95-SRE-KO cells was measured in Transwell migration assays at 48 h. I. The graph depicts the number of invasive HCC95 cells and HCC95-SRE-KO cells shown in Figure S4H. (*P < 0.05, **P < 0.01, ***P < 0.001) J. Western blot analysis of *PTPRZ1* expression in H226 cells transfected with the overexpression plasmid (PTPRZ1-OE), the empty vector (NC) and lip3000 (Con). Each experiment was conducted in triplicate. K. *PTPRZ1* mRNA levels in H226 cells transfected with the overexpression plasmid (PTPRZ1-OE), the empty vector (NC) and lip3000 (Con). Each experiment was conducted in triplicate. L. Western blot analysis of PTPRZ1, c-MYC, Cyclin-D1 and E-cadherin expression in H226 cells and *PTPRZ1*-overexpressing H226 cells. M. The cell migration ability of H226 cells and *PTPRZ1*-overexpressing H226 cells measured in cell scratch assays at 12h and 24h. N. The cell invasion ability of H226 cells and *PTPRZ1*-overexpressing H226 cells measured in Transwell migration assays.

**Figure S5.**
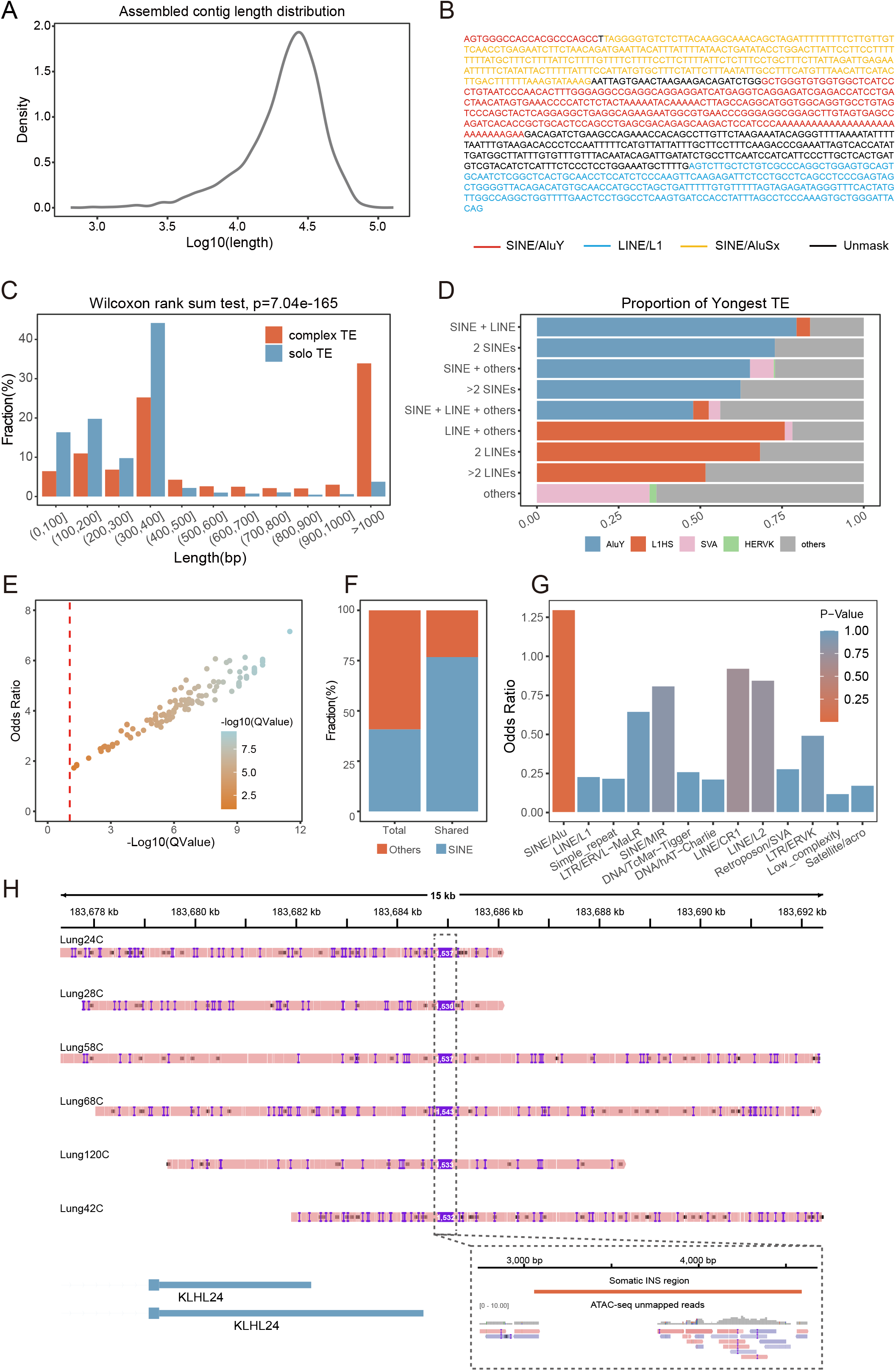
Sequence components and genomic features of complex TEs. A. Distribution of the length of assembled contigs that harbored somatic TEs. B. An example of a complex TE is shown; different colors represent different repeat types. C. A clustered histogram showing the distribution of the lengths of complex TEs and solo TEs. D. Proportion of youngest TE subfamilies in each complex TE component type. E. The proportion of complex TEs in germline and somatic TEs for each sample was used in the statistical analysis. Each dot represents a sample. The colors of the dots and the x-axis indicate the Benjamini‒Hochberg-corrected Q values of Fisher’s exact test. The y-axis represents the odds ratios of Fisher’s exact test. Samples on the right of the red dotted line had significantly different proportions of complex TEs in germline TEs and somatic TEs (Benjamini‒Hochberg-corrected q value < 0.1). F. Proportions of the ratios of total TEs that harbored SINEs and shared complex TEs (shared by at least 2 samples) that harbored SINEs are shown. Orange represents TEs not harboring any SINE segment, and blue represents TEs harboring at least one SINE segment. G. Bar plot showing the enrichment of repeats related to new open chromatin regions compared with genomic repeats annotated with RepeatMasker. All p values were based on one-sided Fisher’s exact test. H. An example of a new open chromatin region related to the insertion of a complex TE. The dashed box indicates the somatic complex TE found at approximately 1537 bp in 6 samples, located approximately 400 bp downstream of *KLHL24*. ATAC-seq unmapped reads were completely mapped to complex TE sequences.

**Figure S6.**
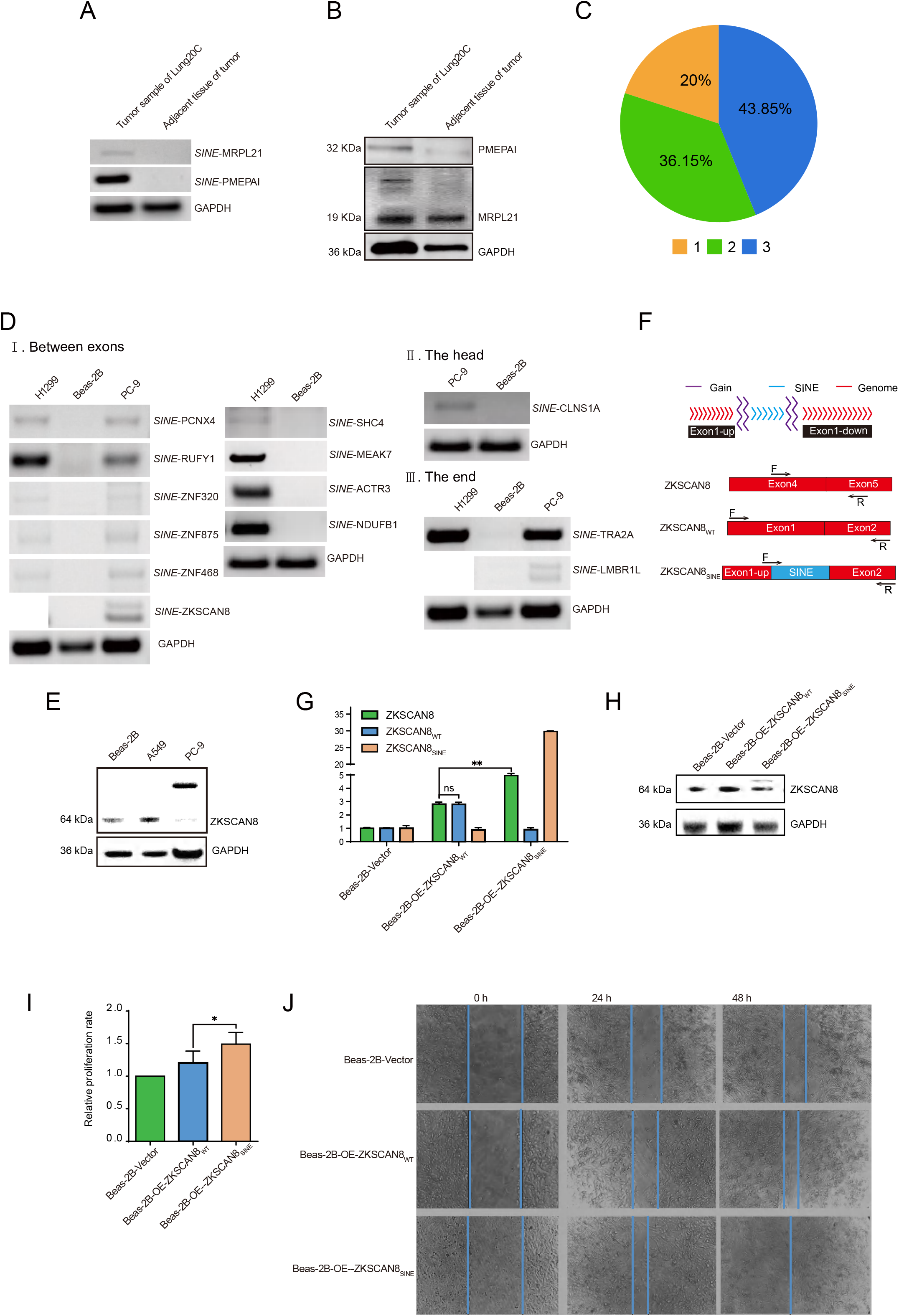
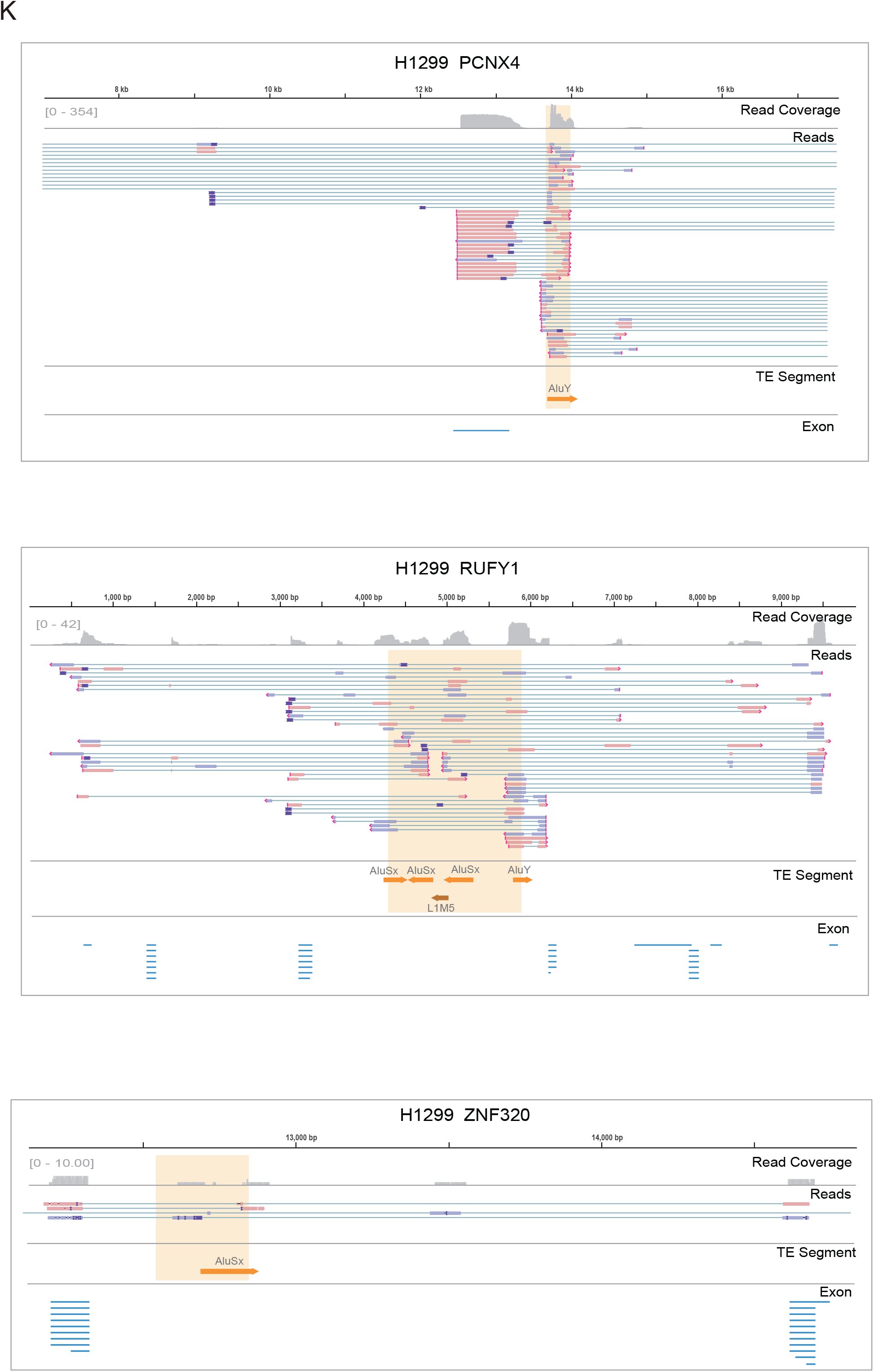
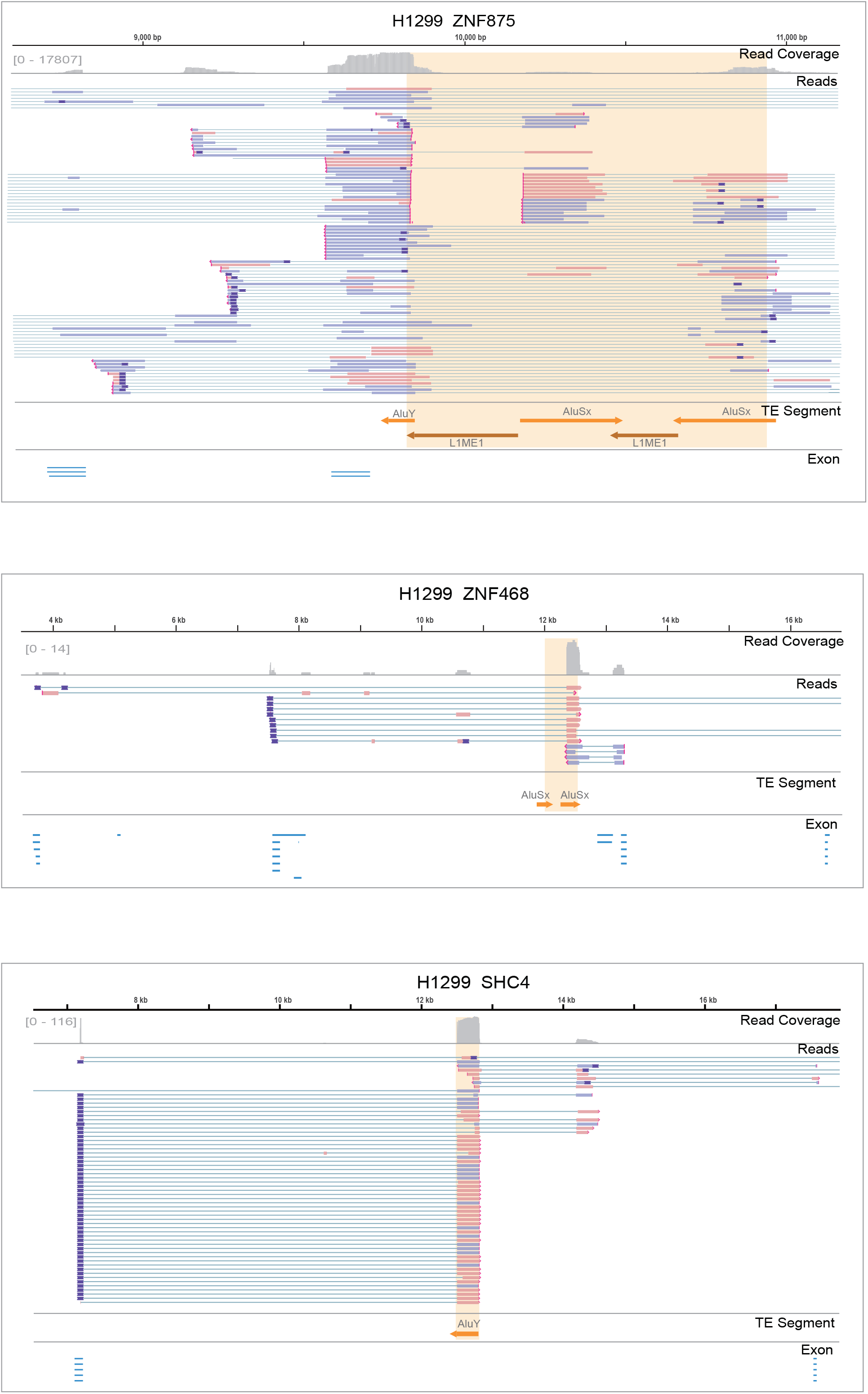
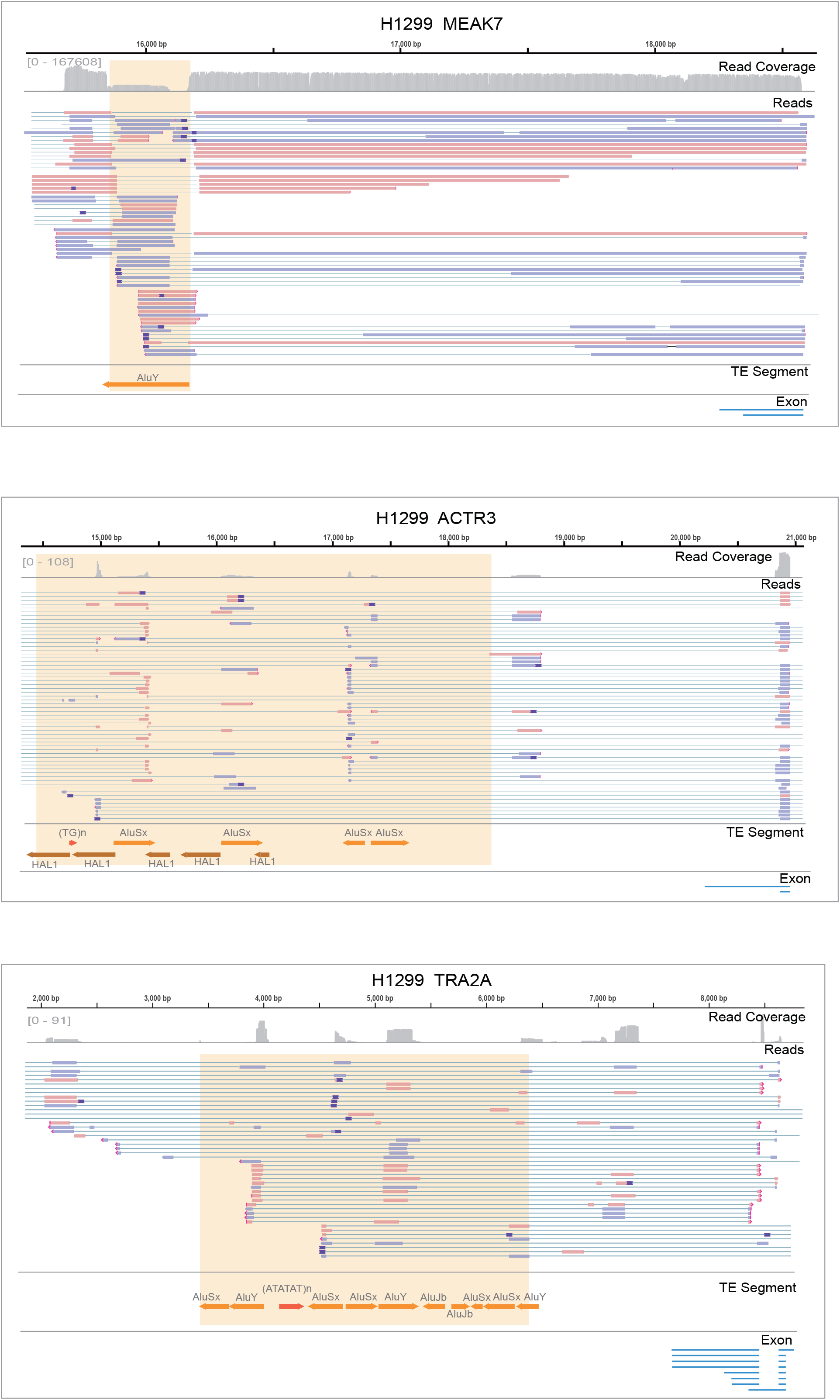
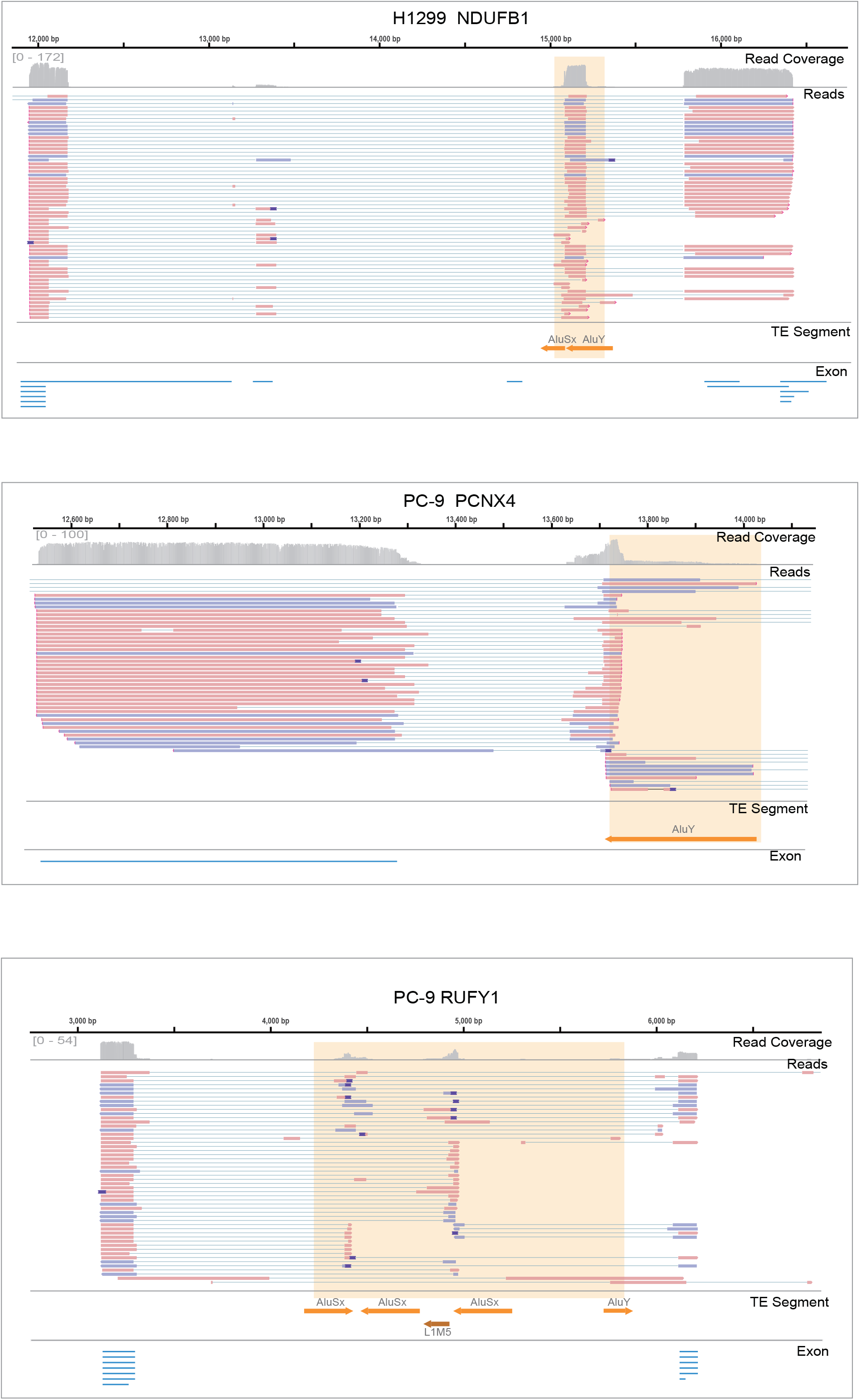
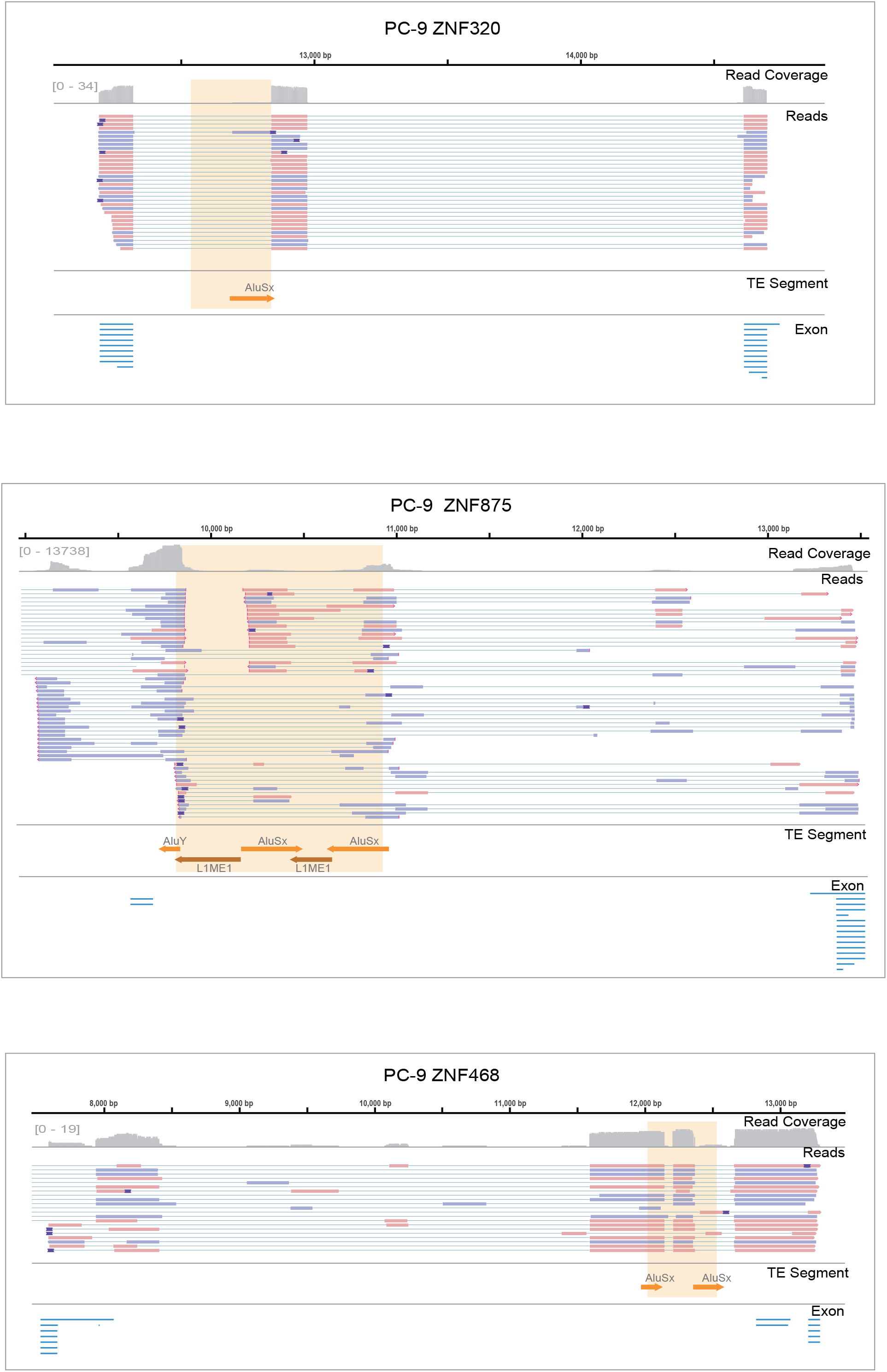
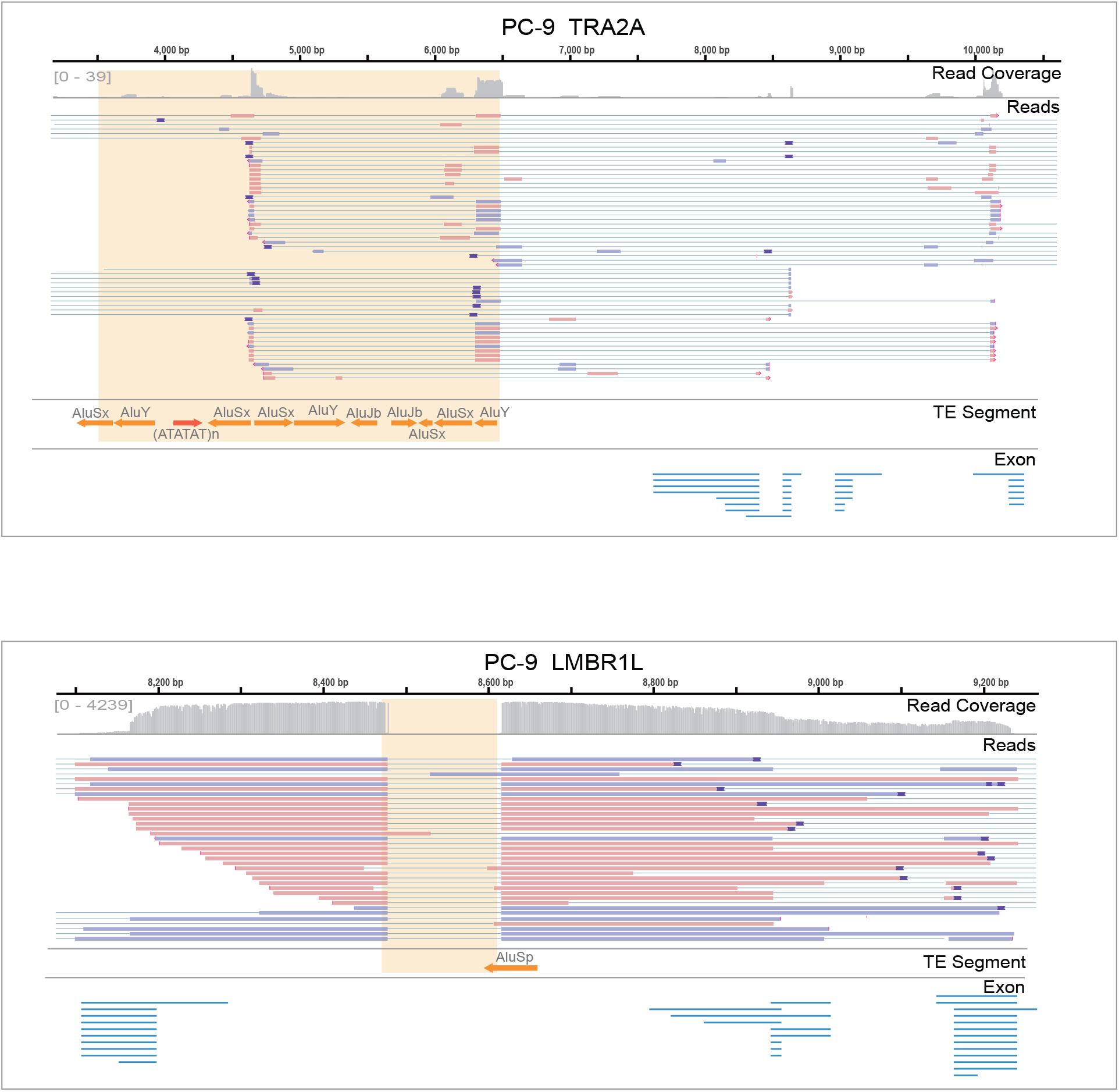
Validation of somatic expressed TEs. A. Agarose gel analysis of SINE expression in tumor samples; somatic expressed SINEs were detected in the *MPRL21* and *PMEPA1* genes. Adjacent tissues of the corresponding patients were used as controls. SINE, short interspersed nuclear element. B. The protein sizes of the two genes involved in S6A were examined by WB. Compared with the proteins expressed in adjacent tissues, SINE expression changed the protein size to varying degrees. C. Pie chart showing the proportion of shared expressed TEs (TE: transposable elements, TD: tandem duplications, *de novo*: *de novo* insertion). D. Agarose gel analysis of SINE expression in PC-9 and H1299 cells is shown. The Beas-2B cell line was used as the control. E. The protein sizes of *ZKSCAN8* in Beas-2B cells. A549 and PC-9 cells were examined by WB. SINE expression changed the size of the protein encoded by *ZKSCAN8* in PC-9 cells. F. The mode of SINE expression in the *ZKSCAN8* gene and the primer positions used to distinguish the regions before and after SINE insertion are shown in the pattern. G. ZKSCAN8_WT_ and ZKSCAN8_SINE_ were overexpressed in the Beas-2B cell line, and the transcript levels of *ZKSCAN8*, ZKSCAN8_WT_ and ZKSCAN8_SINE_ were examined by qPCR. SINE increased the transcription levels of the *ZKSCAN8* gene. H. ZKSCAN8_WT_ and ZKSCAN8_SINE_ were owerexpressed in the Beas-2B cell line and the protein size of ZKSCAN8 were examined by WB. SINE overexpression changed the ZKSCAN8 protein size. I. A CCK8 kit was used to detect cell proliferation after overexpression. J. Representative images of wound-healing assays after overexpression. K. Genome Browser views showing the primary alignments overlapping the sequences of PCR-validated expressed TEs and their adjacent exons. In each panel, the light-yellow shaded area indicates the insertion, while line segments represent the segments of the insertion, different colors represent different types of TE families, and arrows indicate the direction of each TE segment.

## Methods

### Patient Cohort and Ethics

Patients diagnosed with NSCLC were enrolled from West China Hospital of Sichuan University, China. All patients received surgical treatment and did not undergo neoadjuvant therapy before surgery. Tumor tissues and matched blood samples or normal lung tissues were obtained from these NSCLC patients. All diagnoses were verified by histological review by a board-certified pathologist. Cancer clinical stage was defined according to the 8th edition of the American Joint Committee on Cancer (AJCC) TNM staging system. This study protocol was approved by the Institutional Review Board of West China Hospital of Sichuan University (Ethics: Project identification code: 2019.959). All patients provided written informed consent. We collected demographic and clinical information (Table S1).

### Sample Preparation

Blood samples from NSCLC patients were frozen in liquid nitrogen for DNA isolation. Fresh tumor tissues and matched normal lung tissues were divided into two groups: one group of samples was frozen in liquid nitrogen for RNA and DNA isolation, and the other group of samples was digested into single-cell suspensions for ATAC sequencing. The fresh tumor was minced into tiny cubes < 0.5 mm^3^ and transferred into a 15-mL conical tube (BD Falcon) containing 8 mL prewarmed HBSS, 1 mg/mL collagenase I, and 0.5 mg/mL collagenase IV. Tumor pieces were digested on a tube revolver (Thermo Fisher Scientific) for 30 minutes at 37 °C. This suspension was then filtered using a 70-μm nylon mesh (BD Biosciences), and residual cell clumps were discarded. This suspension was rinsed twice with 20 mL PBS and immediately placed on ice. After centrifugation at 500 g at 4 °C for 5 minutes, the supernatant was discarded, and the cell pellet was resuspended in magnetic-activated cell sorting (MACS) buffer (Miltenyi Biotec No. 130-091-222). The tumor cells were purified on MACS columns in MACS buffer. Immune components were depleted using CD45 (Miltenyi Biotec No. 130-045-801) LS columns (Miltenyi Biotec) to obtain a single-cell suspension.

## Method details

### Whole-Genome Sequencing (WGS) and lncRNA Sequencing (RNA-seq)

Genomic DNA was extracted using a DNeasy Blood & Tissue Kit (QIAGEN) according to the manufacturer’s instructions. DNA degradation and contamination were monitored on 1% agarose gels. The DNA concentration was measured with a Qubit® DNA Assay Kit in Qubit® 2.0 Fluorometer (Invitrogen, Grand Island, NY, USA). A total of 0.4 μg DNA per sample was fragmented to an average size of ∼350 bp and subjected to DNA library creation using established Illumina paired-end protocols. A NovaSeq machine (Illumina Inc., San Diego, CA, USA) was utilized for genomic DNA sequencing to generate 150-bp paired-end reads.

RNA extraction was performed with the QIAGEN RNAeasy Mini Kit according to the manufacturer’s instructions. The RNA quality was assessed on a Bioanalyzer 2100 DNA Chip 7500 (Agilent Technologies), and samples with an RNA integrity number (RIN) greater than 7 were further analyzed by RNA-seq. RNA sequencing libraries were generated using rRNA-depleted RNA with the NEBNext® Ultra™ Directional RNA Library Prep Kit for Illumina® (NEB, USA) following the manufacturer’s recommendations. The products were purified (AMPure XP system), and library quality was assessed on the Agilent Bioanalyzer 2100 system. The libraries were sequenced on the NovaSeq machine (Illumina Inc., San Diego, CA, USA), and 150-bp paired-end reads were generated.

### ATAC Sequencing (ATAC-seq)

A total of 50,000 cells were lysed using 50 μL of cold lysis buffer (10 mM Tris-HCl pH 7.4, 10 mM NaCl, 3 mM MgCl2 and 0.1% IGEPAL CA-630). Nuclei were spun at 500 g at 4 °C for 10 minutes. The supernatant was discarded, and nuclei were resuspended in 50 mL reaction buffer containing 5 μL transposase TTE Mix and 10 μL 5×TTBL buffer (Vazyme). The transposition reaction was performed for 30 minutes at 37 °C in a thermomixer shaking at 200 rpm. Directly following transposition, the library was purified using a QIAGEN MinElute Purification Kit (QIAGEN, Hilden, Germany, #28006). Library fragments were amplified using custom PCR primers for a total of 10∼12 cycles. Library quality was assessed on the Agilent Bioanalyzer 2100 system, and 75-bp paired-end sequencing was performed on a NextSeq 500 machine (Illumina Inc., San Diego, CA, USA) to yield on average of 50 million reads per sample.

### PromethION DNA Sequencing

High-molecular-weight (HMW) genomic DNA (gDNA) was extracted from tissues or blood with the MagAttract HMW DNA Kit (QIAGEN). Absolute quantification of the HMW gDNA was performed by using a Qubit dsDNA HS Assay Kit. The DNA library was generated using SQK-LSK109 (Oxford Nanopore Technologies, UK) following the manufacturer’s recommendations. Forty-eight microliters of gDNA (2 μg) plus nuclease-free water (NFW), 3.5 μL of NEBNext FFPE DNA Repair Buffer, 2 μL of NEBNext FFPE DNA Repair Mix, 3.5 μL of NEBNext Ultra II End Prep Reaction Buffer, and 3 μL of NEBNext Ultra II End Prep Enzyme Mix were mixed gently in a PCR tube. After spinning down, the sample was incubated at 20 °C for 5 minutes and 65 °C for 5 minutes, and the mixture was then carefully transferred to a 1.5-mL tube. Sixty microliters of AMPure XP beads was added to the tube mixed by gentle flicking and washed twice with 70% ethanol. Sixty microliters of the end-prepped DNA, 25 μL of Ligation Buffer (LNB), 10 μL of NEBNext Quick T4 DNA Ligase, and 5 μL of Adapter Mix (AMX) were mixed in a new 1.5-mL tube. The sample tube was incubated at 25 °C for 20 minutes. Then, 40 μL of AMPure XP beads was added to the tube, mixing was performed by flicking, and the pellet was resuspended in 25 μL of elution buffer (EB). A total of 30 μL of flush tether (FLT) was added directly to the tube of PromethION Flush Buffer (PFB). Then, 500 μL of Priming Mix was loaded onto the PromethION flow cell for 5 minutes at room temperature. Seventy-five microliters of SQB, 51 μL of LB, and 24 μL of the DNA library (Loading Library) were mixed in 5 minutes, and another 500 μL of Priming Mix and 150 μL of Loading Library were loaded onto the flow cells. Then, the DNA library was sequenced on a PromethION machine (Oxford Nanopore Technologies Inc., Oxford, England). Reads were base called in batches by Guppy 3.2.8 using the default parameters during sequencing.

### Construction and sequencing of Hi-C libraries

Hi-C libraries were created from one tumor sample and its paired blood sample as described previously [46]. Briefly, the samples were fixed with formaldehyde and lysed, and then DNA was cross-linked with DNA or intracellular proteins. Cross-linked DNA was digested with DpnII restriction enzyme to produce sticky ends on both sides of the cross-linking. Sticky ends were labeled using biotinylated nucleotides and then proximity ligated to form chimeric junctions. After reversal of crosslinks, ligated DNA was purified and then broken into 300-to 700-bp fragments. The DNA fragments containing the interaction were pulled down with streptavidin beads to construct the Hi-C library. The concentration of the library and the size of the inserted fragment (insert Size) were determined with Qubit 2.0 and Agilent 2100, respectively. The effective concentration of the library was quantified accurately by qPCR. Paired-end reads (150 bp) were produced on a NovaSeq machine (Illumina Inc., San Diego, CA, USA).

### Data Processing for Next-Generation Whole-Genome Sequencing

The clean paired-end reads were aligned to the hg38 reference genome using Burrows‒ Wheeler Aligner (BWA) MEM (version 0.7.17) [75]. The mapped reads were converted into sorted and indexed BAM format files using SAMtools (version 1.9) [76]. Duplicate reads were marked using the MarkDuplicates module of the Genome Analysis Toolkit (GATK [77], version 4.1.2.0). To improve the quality of somatic indel identification, we performed local realignment and base quality score recalibration using GATK BaseRecalibrator and ApplyBQSR modules. Finally, somatic SNVs and indels were called from tumor and matched blood sample pairs using Mutect2 from GATK, and the variants detected by Mutect2 were filtered with GATK FilterMutectCalls. We analyzed mutational signatures by linear decomposition using Mutalisk (Lee et al., 2018). Mutation significance analysis was carried out using the MutSig2CV algorithm, as elaborated in several genome projects [78]. Copy number variants were identified using Control FreeC (version 11.6) [79] with the default parameters. Tumor purity was estimated using 2 tools: we used CNV Facets [80] with the parameters “--snp-nprocs 30 --cval 25 400 --nbhd-snp 500” to estimate tumor purity; we used ABSOLUTE [81] with the parameters “ploidy = 2 breakPointThreshold =.8 window = 50000” to estimate tumor purity.

### RNA-seq Data Analysis

For each sample, paired-end reads were aligned to the hg38 reference genome using HISAT2 (version 2.1.0) [82, 83]. SAMtools was used to sort and index the bam files. StringTie (version 1.3.6) [84] was used to generate the GTF files containing the read count information and calculate the TPM matrix of gene expression for all genes across all samples. Only genes with a mean TPM greater than 1.0 in all samples were considered expressed genes.

### ATAC-seq Data Analysis

Adapters of raw paired-end ATAC-seq reads were trimmed using Cutadapt (version 1.15) [85]. The adapter-trimmed paired-end reads were aligned to the hg38 reference genome via BWA with the ‘mem -M -t 10 -R’ options. SAMtools was used to sort and index the BAM files. Duplicated reads were subsequently marked and removed using the GATK MarkDuplicates tool. To generate a high-quality mapping profile for postprocessing, we removed the following reads: MAPQ (mapping quality) < 30, not paired-aligned, mapped to ChrM, and TLEN (observed template length) > 2,000 bp. Then, MACS2 (version 2.2.7) [86] was used to call peaks from the filtered BAM files with the following parameters: -f BED --nomodel --nolambda --shift 100 --extsize 200 -B --call-summits --SPMR. Finally, for downstream analysis, we constructed a consensus open peak set using peaks from all samples, as suggested in a previous study (Corces et al., 2018).

### Oxford Nanopore Sequencing Data Processing

ONT long reads were aligned to the hg38 reference genome using NGMLR (version 0.2.8) [31] with the following parameters: -t 40 -x ont. A set of structural variations (SV) was obtained using Sniffles (version 1.0.11) [31] for each genome in a sensitive fashion (using -s 5 –genotype –cluster -q 20 -n −1). All supporting reads were reported per variant in the variant call format (VCF) file. SVs were discarded when all the supporting reads were less than 1000 bp. Other optional parameters were left as default. Insertions, deletions, and duplications shorter than 30 bp were discarded.

### Hi-C Data Analysis

Clean reads were aligned to the hg38 reference genome using BWA aln (version 0.7.10-r789) with the default parameters. Then, HiC-Pro (version 2.10.0) [87] was used to analyze the alignments and identify the valid/invalid interaction pairs. The identification of topology-related domains (TADs) was performed according to genomic bins of 40 kb using TadLib. HOMER [88] was used to calculate the inclusion ratio (IR). TADs with IR > 1 were retained, and TADs with a length of less than 5 bins were filtered out.

### Somatic SV Identification

We used an in-house script, which used the supporting split reads of insertions (INSs) and duplications (DUPs) reported by Sniffles, to detect somatic insertions. INSs and DUPs of tumor samples identified by Sniffles with a length of less than 50 bp were discarded before further analysis.

The initial detection of somatic INSs and DUPs follows the same principle. Here, we take somatic INSs as an example to describe the steps of initial somatic INS identification. First, we grouped the distances between every two adjacent INSs in a tumor sample and sorted the distances in ascending order. If the distance between two adjacent INSs was less than the 10th percentile of the distance group, we determined that these two INSs were the same INS. Second, for each INS, we chose the region between the breakpoint supported by the highest number of alignments and the breakpoint identified by Sniffles as the candidate break region. Third, we removed INSs when there was no significant difference (Kolmogorov‒Smirnov test, p value < 0.05) between the counts of split alignments in the candidate break regions in the tumor and its matched blood sample. Fourth, we defined the candidate region as the region between the left candidate break region and the right candidate break region. We removed INSs when there was no significant difference (Wilcoxon rank-sum test, p value < 0.05) between the size of the candidate regions in the tumor and its matched blood sample. Fifth, INSs that were uniquely identified in the tumor sample were retained for the next filter. Finally, we manually reviewed the somatic INSs using the Integrative Genomics Viewer based on the split alignments of INSs that passed the above filters, and INSs exclusively found in the tumor sample remained.

To prove the validity of our somatic INSs, we collected all available long-read whole-genome sequencing (WGS) data (generated on the ONT platform) from the 1000 Genomes Project [37] and identified insertions and duplications using Sniffles. We also downloaded a germline SV dataset (generated on the ONT platform) from 405 unrelated healthy Chinese individuals [89]. The SVs from the two projects were pooled together to be employed as an ONT platform-based ‘Panel of Normals (PONs)’. Specifically, the structural variation comparison tool Jasmine [] was used to merge SVs from these datasets using the following parameters: --dup_to_ins --ignore_strand max_dist=100 min_seq_id=0.9. Then, the same parameters were used to compare the somatic INSs we identified with the germline INSs in PONs.

### Local Assembly of Somatic Insertions

After somatic INS identification, we attempted to locally assemble all somatic INSs through the following process to classify INSs into somatic TDs, somatic TEs and *de novo* insertions. First, the INS-supported reads for each somatic INS were extracted based on the record of Sniffles and were assembled using Shasta (version 0.5.1) [32] with the following parameters:

- Reads.minReadLength (1000)
- MinHash.minHashIterationCount (60)
- MinHash.minFrequency (2)
- Align.minAlignedMarkerCount (100)
- MarkerGraph.minCoverage (5)

To further improve the quality of the assembled Shasta sequences, Racon (version 1.4.3) [33] and Medaka (version 1.0.3, https://github.com/nanoporetech/medaka) were used to polish the draft assemblies. For draft assembly of each somatic INS, INS-supported reads were used for iteratively running 4 rounds of Racon. After the initial polishing with Racon, further polishing was performed using the r941_prom_high_g360 model in Medaka. Finally, the polished assembled sequences were aligned against the hg38 reference genome via minimap2 (version 2.17) [67] for downstream analyses.

### Somatic Insertion Annotation

Assembled somatic insertion sequences with flanking 50-bp reference sequences were annotated by RepeatMasker [35] with additional annotation by Tandem Repeat Finder (TRF) [35] and Sdust [36], as polymorphic tandem repeats were excluded in the RepeatMasker result. Any region on assembled insertion sequences with no transposon annotation from RepeatMasker was reannotated with TRF. We classified TRF annotation results based on the length of the repeat pattern into simple repeats (≤10 bp), tandem repeats (> 10 bp and <= 100 bp), and satellites (> 100 bp). In addition, assembled insertion sequences marked as “Unmask” by RepeatMasker were annotated by TRF and Sdust if the annotation length was greater than 5 bp.

### Somatic Insertion Classification

Assembled somatic insertion sequences were classified as somatic TDs, somatic TEs, and somatic *de novo* insertions by comparison with inserted site or duplicate-starting site adjacent reference genome sequences in two ways. In the whole segment method, upstream and downstream reference sequences with a length of 50 bp longer than the insertion sequences were extracted, and the similarity value of the adjacent sequences to the insertion/duplication sequences was calculated as described by Needle [90] (EMBOSS, version 6.6.0.0). Insertion sequences with identity to adjacent reference sequences greater than 90% were considered TDs. In the repeat pattern method, insertion sequences were annotated by TRF first to identify repeat patterns. For insertion sequences that had greater than 90% of the region annotated by TRF and only one repeat pattern, upstream and downstream reference sequences with a length of 1.5-fold the repeat pattern plus 10 bp were extracted. The repeat pattern was compared with adjacent reference sequences by pairwise local alignment (Gotoh local alignment [91]). Insertion sequences with repeat patterns that had a similarity score greater than 85% with adjacent reference sequences were considered TDs. The insertion sequences except TDs that harbored any TE segments were classified as TEs. All insertion sequences other than TDs or TEs were classified as *de novo* insertions.

### Recurrent Insertion Discovery

Recurrent INS discovery was carried out separately for TEs, TDs and *de novo* INSs in LUSC samples and LUAD samples using the R package fishHook [38]. The hypothesis interval was set as a 1 kb genomic window. Heterochromatin and active chromatin states of genomic regions were extracted from the chromHMM data of the A549 cell line from Epigenomics Roadmap (Roadmap Epigenomics Consortium et al., 2015) and added to fishHook as covariates. Any fishHook outcome with an FDR < 0.05 that was shared by at least three samples was regarded as recurrent.

### LUSC-enriched somatic SRE Identification in WGS SRS Data

The LUSC-enriched somatic SRE was identified using ExpansionHunter Denovo (version 0.9.0) with the following parameters:

- min-anchor-mapq (50)
- max-irr-mapq (40)

Any outcomes with at least one read supporting the repeat pattern “AAAG” expansion at chr7: 121602900-121604000 were deemed the LUSC-enriched somatic SRE.

### Eligible Region Definition

The hg38 genome was divided into 500 bp windows, and the genomic coverage in each window was measured by mosdepth (version 0.3.3) (Pedersen et al., 2018). For each chromosome in each sample, the 25th (Q_25_) and 75th quantiles (Q_75_) were computed. The lowest coverage threshold, L_cov_, was determined for each chromosome based on the following formula:

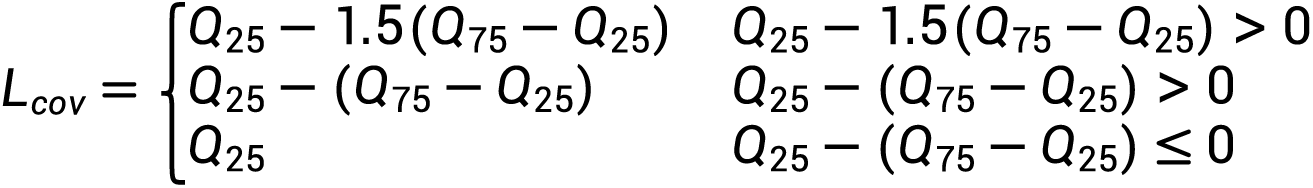

Genomic windows with coverage above the threshold were considered valid; otherwise, they were considered invalid. Thus, the validity of every genomic window for each sample was determined. To be considered eligible, a genomic window had to be valid in more than 90% of the samples.

### Validation of the LUSC-enriched Somatic SRE in Tissues and Cell Lines using Targeted Sequencing

DNA extraction and PCR: DNA was extracted from tumor tissues, normal tissues, LUSC cell lines (H520, KNS62, HCC95, CALU-1, H1703, H226, and SK-MES-1), LUAD cell lines (PC-9 and A549), U251, BEAS-2B and large cell lung cancer H1581 cells using the DNeasy Blood & Tissue Kit (QIAGEN) according to the manufacturer’s instructions. The forward and reverse primers were designed according to the 1 kb upstream and downstream sequences of the insertion position and synthesized by Sangon Biotech. DNA was extracted from clinical samples or cell lines and subjected to PCR using primers. Each 50-μL reaction consisted of 2× PremSTAR Max Premix 25 μL, 0.8 μL 10 μM anchor primer, 0.8 μL 10 μM test primer, and 150 ng DNA. PCR was performed as follows: an initial step of 2 minutes at 98 °C, 30 cycles of 10 seconds at 98 °C, 15 seconds at 65 °C and 60 seconds at 72 °C (size of PCR products between 1 kb and 1.5 kb). Finally, the samples were incubated at 72 °C for 2 minutes. The PCR product was purified by a column purification system, and the concentration of purified DNA was detected by a Qubit kit. The concentration of purified DNA was required to be 10 ng/μL, and the volume was at least 20 μL. The size distribution of the DNA fragments was determined by LabChip GXII Touch HT (PerkinElmer).

#### Library construction and sequencing

A DNA library was generated using a Barcode kit (QitanTech, CN) following the manufacturer’s recommendations. A total of 0.2 pmol of cleaned PCR products per sample with 7 μL of DNA repair buffer, 3 μL of end-prep mix and nuclease-free water (up to 60 μL) were used in the end-repair steps: 10 minutes at 20 °C, 10 minutes at 65 °C and hold at 4 °C. Then, the reactions were purified with a 1.0 × volume of VAHTS DNA Clean Beads and eluted with 35 μL of nuclease-free water. Barcodes supplied in the kit were attached to the PCR product ends, 5 μL (1 pmol) of barcodes, 15 μL 4x ligation reaction buffer and 5 μL of DNA ligase enzymes were added to each end-prepped PCR product, followed by incubation for 10 minutes at room temperature. Then, the reactions were purified with a 0.8 × volume of VAHTS DNA Clean Beads and eluted with 15 μL of nuclease-free water. Equimolar amounts of each barcoded sample were pooled in a 1.5 mL Eppendorf DNA LoBind tube, ensuring that sufficient sample amounts were combined to produce a pooled sample of 0.3 pmol total. To accomplish the ligation, 5 μL of Sequencing Adapter Complex (SAC), 25 μL of 4x ligation reaction buffer (LRB), and 10 μL of DNA ligase enzymes were added to the product, followed by incubation for 10 minutes at room temperature. After ligation, each library was purified with a 0.4 × volume of VAHTS DNA Clean Beads, and short-ligation wash buffer (SWB) was used to enrich fragments > 200 bp. After removing the liquid, 15 μL of AE elution buffer (AEB) was used to collect the libraries, and the libraries were quantified using a dsDNA HS Kit on a Qubit fluorometer. A total of 8 μL of Priming Anchor (PRA) was added to 392 μL of priming buffer (PRB). Then, 200 μL of the mixture was loaded onto the Qcell-384 flow cell for 15 minutes at room temperature. Then, 100 μL of sequencing reaction buffer (SRB), 40 fmol library, and nuclease-free water (up to 200 μL) were mixed (library mix), and 200 μL of the library mix was loaded onto the flow cells. Then, the DNA library was sequenced on a QNome-3841 device. Kirky@v2 (QitanTech) was employed for base calling.

#### Data analysis

Raw data in fastq format were first processed through NanoFilt [92]. In this step, clean data were obtained by removing reads whose sequencing quality was less than 7 and length was less than 500 bp. Clean reads were then aligned against the hg38 reference genome using NGMLR (version 0.2.8) with the following parameters: -t 20 -x ont. To identify the LUSC-enriched somatic SRE, first, alignments in chr7:121603000-121604000 were extracted. Then, reads harboring an SRE similar to the LUSC-enriched somatic SRE were detected by an in-house script. This script used TRF to identify the repeat pattern of the SRE. The ratio of reads harboring the SRE, which is similar to the LUSC-enriched somatic SRE, to reads aligned into chr7:121603000-121604000 was calculated. The tumor sample was considered to harbor an LUSC-enriched somatic SRE when the ratio was greater than 0.6% and the ratio of its matched blood sample was less than 0.6%.

### Cell Culture

Human cell lines provided by the American Type Culture Collection (Manassas, VA, USA) were used in the present study. The cells were incubated in Dulbecco’s modified Eagle’s medium (DMEM) or RPMI 1640 medium (Gibco; Thermo Fisher Scientific, Inc.). The medium was supplemented with 10% fetal bovine serum (FBS) (Gibco; Thermo Fisher Scientific, Inc.), and the cells were all maintained at 37 °C in 5% CO_2_. Upon reaching 70-80% confluence, the cells were washed with PBS and detached with 0.25% trypsin/0.2% ethylenediaminetetraacetic acid (EDTA). Cell morphology was viewed under a light microscope, and the cells were suspended to a concentration of 1 x 10^6^ cells/ml.

### Real-Time Quantitative PCR

Total cell RNA was extracted from LUSC cells (H520, HCC95, KNS62, H1703, H226, CALU-1 and SK-MES-1), LUAD cells (PC-9 and A549), U251, BESA-2B and H1581 cells, using the RNeasy Mini Kit (QIAGEN). RNA quality was verified by agarose gel electrophoresis and converted into cDNA using the iScript cDNA Synthesis Kit (Bio-Rad). SYBR Green Supermix (Bio-Rad) was used for real-time quantitative PCR. The reaction conditions and PCR system were operated in accordance with the instructions. All sequences were designed and synthesized by TSINGKE (Chengdu, China) and are listed in the oligonucleotide table in the Key Resources section (SRE forward, ATGCGAATCCTAAAGCGTTTCCTCG; SRE reverse, AACTAAAGACTCTAAGCTCTCAGC). Target mRNA levels were measured using the 2^-ΔΔCt^ method.

### Western Blotting

The total protein contents of the U251, H520, HCC95, KNS62, H226 and H1581 cell lysates were quantified by using the bicinchoninic acid (BCA) assay, subjected to sodium dodecyl sulfate‒polyacrylamide gel electrophoresis (SDS‒PAGE) (Beyotime, China) and electrotransferred to polyvinylidene difluoride (PVDF) membranes (Millipore, USA). After blocking with 5% bovine serum albumin Tris-buffered saline-Tween (BSA TBS-T) solution, the membranes were incubated with primary antibodies against *PTPRZ1* (1/1000) (Abcam, USA). The protein was visualized by chemiluminescence with an enhanced chemiluminescence (ECL) kit (Beyotime, China) and quantified by densitometry, which was normalized to the housekeeping protein glyceraldehyde 3-phosphate dehydrogenase (GAPDH) or β-actin.

### Immunofluorescence Staining

Slides with methanol-fixed cells and 0.5% Triton X-100-transparent cells were placed for 20 minutes at room temperature and subsequently incubated first with 10% goat blocking serum for 1 hour and then with diluted primary antibodies at 4 °C overnight. The slides were further incubated with fluorophore-conjugated secondary antibody for 2 hours. The cell nuclei were counterstained with 4,6-diamino-2-phenyl indole (1/1000, DAPI) for 5 minutes. Images were obtained using a Leica Stellaris spectral confocal microscope. The primary antibodies were as follows: anti-*PTPRZ1* (1/300, Abcam, USA). To determine *PTPRZ1* expression in cells, fluorescence intensity was measured. For each fluorescence intensity, *PTPRZ1* expression was calculated in five randomly selected microscopic fields with 40 cells at 400X by Image software. All immunofluorescence staining was independently repeated at least three times.

### Knockout of the LUSC-Enriched Somatic SRE by CRISPR/Cas9 in LUSC cell lines

The somatic SRE in HCC95 cells (SRE long insertion) was knocked out by CRISPR‒Cas9-containing expression cassettes for hSpCas9 and the chimeric guide RNA. To target the SRE in HCC95 cells, two guide RNA sequences of GACCACTGGGAAAATGCAGA and TTAAATGCAGAATATCTGAA were selected via the http://crispr.mit.edu website, and two guide RNA sequences of CATAATCATGGTGCCATGAT and TTAAATGCAGAATATCTGAA were selected to target SRE in HCC95 cells. Plasmids containing the guide RNA sequence were transfected into HCC95 cells using a Lipofectamine® 3000 Transfection Kit according to the manufacturer’s instructions (Thermo Fisher Scientific).

### Viral transduction

For shRNA knockdowns, virus particles were produced by co-transfection of 293T cells with pLVX /puro construct expressing the indicated shRNA, psPAX2 and pCMV-VSVg in a ratio of 5:3:2 by mass. The *PTPRZ1*-shNRA oligo sequences: GCTGCTTTAGATCCATTCATA. After 48 h of transfection, HCC95 cells were transduced with 0.45-μm filtered viral supernatant and 8 μg/ml polybrene. Cells were selected 24 h after medium replacement with 2 μg/ml puromycin.

### Cell Lipid Transfection

H226 cells (*PTPRZ1* low expression) were seeded on coverslips in 6-well plates at a density of 40 × 10^4^ cells per well and cultured for at least 24 hours before transfection to attain 80–90% monolayer confluence. Then, the samples were washed with sterile PBS, and 500 µL of serum-free 1640 medium containing 2 µg of the *PTPRZ1*-pcDNA3.1 overexpression plasmid, empty vector pcDNA3.1 and lip3000 (Invitrogen) was added to each well. The wells were incubated without serum for 4 hours. Then, the samples were washed with PBS, and 1.5 mL of complete 1640 medium was added to each well. Forty-eight hours after transfection for Western blotting assay.

For ZKSCAN8_WT_ and ZKSCAN8_SINE_ insertion plasmid construction, the ZKSCAN8_WT_ and ZKSCAN8_SINE_ fragments were amplified by PCR using the complementary DNA (cDNA) of Eeas-2B and PC-9 cells as templates. Then, the amplified fragment was inserted into the pcDNA3.1-EGFP vector, and the constructed positive plasmid was confirmed by DNA sequencing. Recombinant plasmids with the ZKSCAN8_WT_ or ZKSCAN8_SINE_ sequence were generated by transfection in Beas2B cells as described above.

### Wound Healing Scratch Assay

The migration of cells was evaluated with a wound healing scratch assay. H226 cells were seeded in 6-well plates (1 × 10^6^/well) in culture medium and cultured until confluent. Then, the cells were treated with *PTPRZ1*-pcDNA3.1, pcDNA3.1 and lip3000. Then, an artificial scratch wound was drawn at the center of the well and photographed. After 24 hours of culture, the cell scratch wound was photographed again, and the migration distance was determined by the ratio of healing width at 24 hours vs. the wound width at 0 and 12 hours. Each assay was performed in triplicate and repeated three times.

The wound healing scratch assay was also carried out in Beas-2B cells after the transfection of the recombinant plasmids with the ZKSCAN8_WT_ or ZKSCAN8_SINE_ sequence for 24 h. The pcDNA3.1-EGFP vector was also transfected as a control.

### Cell Counting Kit 8 (CCK-8) Assay

The Cell Counting Kit 8 (CCK-8) Assay was performed to test cell viability. In brief, HCC95 cells and PTPRZ1-shRNA cells were seeded in 96-well plates, and each well was seeded with 5000 cells. Then, the cells were cultured in a 37°C, 5% CO_2_ incubator for 24 h, 48 h, 72 h, 96 h, and then 10 μL CCK-8 reagent was added to each well in an incubator for 1 h. The absorbance was determined at 450 nm by a microplate reader (BioTek Epoch2). Each group was tested three times.

### IncuCyte cell proliferation assays

IncuCyte cell proliferation assays were performed to test cell proliferate ability. In brief, 5000 cells were seeded per well in corning 96-well plates, and then cells were cultured in an incubator with an IncuCyte microscopy-platform (Sartorius, IncuCyte S3). Phase-contrast images were acquired every 6 hours for 2-3 days until cells reached >80% confluence. Cell proliferation was monitored as the occupied area (% confluence) of cell images over time and calculated as fold of cell confluence normalized to 0h. Cell confluency percentages were calculated using the integrated Cell Player software and results were analysed with Prism 8 (GraphPad Software).

### MTT Assay

The 3-(4,5-dimethyl-2-thiazolyl)-2,5-diphenyl-2-H-tetrazolium bromide (MTT) assay was performed to test cell viability. An MTT cell proliferation and cytotoxicity assay kit (Beyotime Biotechnology, Shanghai, China) was used in the present study. In brief, hippocampal neurons (7 DIV) in 96-well plates were incubated with 100 μl MTT (0.5 mg/ml, dissolved in NF media) for 4 h, and 100 μl dimethyl sulfoxide (DMSO) was added to dissolve the dark blue formazan crystals. The absorbance was determined at 570 nm by a microplate reader. Finally, the cell viability rate was expressed as a ratio (%) between the absorbance of the treated group and that of the control group.

### Transwell Migration and Invasion Assay

To perform Transwell migration and invasion assays, H226 cells were cultured with tumor supernatant in 24-well plates and reached 90% confluence after transfection. Then, the samples were washed with sterile PBS, and 500 µL serum-free RPMI 1640 containing 1 µg *PTPRZ1*-pcDNA3.1 and pcDNA3.1 was added to each well for 24 hours. H226 cells were cultured with tumor supernatant (FBS free) in 24-well plates. Transwell migration assays were performed in 24-well plates with an 8-lm pore polycarbonate membrane (Corning, Tewksbury, MA). The cells (3.5 × 10^4^) were starved for 12 hours and then added to the upper chambers of the wells in 0.5 mL of serum-free medium. A total of 600 μL of Roswell Park Memorial Institute (RPMI)-1640 medium supplemented with 10% fetal bovine serum was added to the lower chamber. The cells were incubated for 24 hours at 37 °C to enable cell invasion into the lower chamber. The invaded cells were fixed and stained with 0.5% crystal violet. Images were taken by microscopy (Olympus, Tokyo, Japan) in at least eight representative fields. The invasion assay was performed with filters precoated with Matrigel (BD Biosciences, Franklin Lakes, NJ), and the following protocols were the same as those described for the migration assay.

### Dual-Luciferase Reporter Assay

Two enhancer sequences (enhancer1 and enhancer2) flanking the SER insertion site, the promoter sequence *of PTPRZ1*, and the predicted CTCF binding site sequence were cloned into the pGL3 basic vector, and CTCF genes were cloned into the pcDNA3.1 basic vector. The constructed positive plasmid was confirmed by DNA sequencing. HEK293T cells were grown in 6-well plates to 80% confluence before transfection with reporter plasmids. Cells were cotransfected with 200 ng of enhancer1, enhancer2 or PTPRZ1 promoter-luciferase reporter constructs, 10 ng of Renilla luciferase plasmid (pRL-TK), pcDNA3.1-CTCF or control plasmid, followed by incubation for 24-48 hours. Firefly luciferase activities were determined using a dual-luciferase reporter assay system (Promega) according to the manufacturer’s protocol, and relative luciferase activity was determined by normalizing firefly luciferase activity against Renilla luciferase activity. All experiments were performed in triplicate.

### Immunohistochemistry Assays

Formalin-fixed, paraffin-embedded (FFPE) blocks of LUSC were used for immunohistochemistry staining based on standard staining methods, including a deparaffinization and rehydration step (Autostainer XL Leica ST5010), quenching in endogenous peroxidase blocking buffer (3% H_2_O_2_) for 15 minutes at room temperature (RT), an acidic antigen retrieval step (heating for 8 minutes twice at 100 °C in citric acid pH 6), and blocking with 10% goat serum. A primary antibody (rabbit anti-*PTPRZ1* polyclonal antibody, Absin#abs134714, 1:200) was applied to slides overnight at 4 °C, and a secondary antibody (peroxidase-conjugated AffiniPure goat anti-rabbit IgG (H+L), JIR# 111-035-003, 1:250) was added for 40 minutes at 37 °C. Then, the location of peroxidase activity was stained with a One Drop SignalStain® DAB Substrate Kit (Cell Signaling, #8059). Cell nuclei were counterstained with hematoxylin. Slides were scanned using the Nikon DS-Ri2 automated slide scanner at 20X.

### 3C Library Generation

Here, 3C experiments were performed in HCC-95 and H226 cell lines. Formaldehyde crosslinking and 3C library preparation were performed as described previously[93]. Briefly, 1 × 10^7^ were cross-linked with formaldehyde for 10 minutes and quenched with a final concentration of 0.125 mM glycine for 5 minutes at room temperature. Cells were counted and lysed with 500 μL 1× cold lysis buffer (10 mM Tris-HCl pH = 8.0, 10 mM NaCl, 0.2% Igecal CA630) containing 1× protease inhibitor (Roche, Indianapolis, IN, USA) for at least 15 minutes on ice. Cell nuclei were pelleted, washed with 500 μL ice-cold 1× MseI buffer (NEB, Ipswich, MA, USA), resuspended in 500 μL 1× MseI buffer with 0.3% SDS, followed by incubation for 1 hour at 37 °C. Then, 1% Triton X-100 was added to the sample, and the sample was incubated for another 1 hour to sequester the SDS. Each sample was digested overnight with 600 U restriction enzymes at 37 °C. To stop the restriction digestion, 1.6% SDS (final concentration) was added, and the samples were incubated at 65 °C for 20 minutes. Ligation was performed at 16 °C for 4 hours in 15-mL tubes containing 745 μL 10× T4 ligase Buffer, 10% Triton-X 100, 80 μL 10 mg/mL BSA, 6 mL water, 575 μL of cell lysate, and 10 μL 1 U/μL T4 ligase (Invitrogen). The crosslinks were reversed with Proteinase K (Invitrogen) at 65 °C overnight. 3C samples were then purified using phenol chloroform extraction and quantified by Qubit dsDNA HS Assay (Life Technologies).

### PCR Primers and 3C-PCR

Primers were designed to detect the interaction between *PTPRZ1* and the SRE-inserted enhancer regions. The anchor primers were designed near *PTPRZ1*. The test primer was designed around the MseI cutting sites near the regions of enhancer 1 and enhancer 2. Each test primer was paired with the anchor primer. The sequences of the primers are listed in Supplementary Table S6A. All PCRs were performed with PremSTAR Max Premix (2x) (Takara R045). Each 50-μL reaction consisted of 25 μL of 2× PremSTAR Max Premix, 0.8 μL of 10 μM anchor primer, 0.8 μL of 10 μM test primer, and 100 ng of 3C DNA. PCR was performed as follows: an initial step of 2 minutes at 98 °C, 30 cycles of 10 seconds at 98 °C, 15 seconds at 60 °C and 60 seconds at 72 °C. Finally, the samples were incubated at 72 °C for 2 minutes. The products were purified (Invitrogen) and sequenced (Tsingke, Beijing, China).

### Detection of New Open Chromatin Regions Related to Somatic Complex TEs

For each tumor sample, three types of reads, split reads and discordant reads within 500 bp of the TE insertion site, and all unmapped reads, were extracted by SAMtools and the Python module pysam based on the FLAG field from the ATAC-seq BAM file. The hg38 reference genome and all assembled somatic complex TE contigs were pooled together as the new reference genome. Then, extracted reads were aligned to the new reference genome using BWA. Pysam was used to extract reads properly mapped to the assembled complex TE sequence. When more than one pair of reads mapped to the complex TE, the TE was considered to be associated with a new open chromatin region.

### Detection of Somatic TE Expression

Short-read RNA-seq data of our cohort: Unmapped reads, split reads, and discordant read pairs were extracted by using SAMtools and the Python module pysam based on the FLAG field from the RNA-seq BAM files. For each tumor sample, the extracted RNA-seq reads were aligned to each assembled somatic insertion contig, which harbored TE segments, using HISAT2. To identify expressed TEs, gene annotation of GENCODE (version 33) was projected onto the assembled somatic insertion sequences. The expressed TEs were determined as reads aligning to the inserted region or spanning the breakpoint between the inserted region and the adjacent exon as well as read pairs mapped to both the inserted region and the adjacent exon.

Long-read RNA-seq data of cell lines: Long-read RNA-seq data of 3 cell lines were downloaded from the DDBJ database. To increase the amount of sequencing data for A549 cells, we generated nearly 6 GB of long-read RNA-seq data. The final amounts of long-read RNA-seq data for the 3 cell lines were 8.25 Gb for H1299 (N50 = 2.569 kb), 5.96 Gb for PC-9 (N50 = 1.949 kb), and 8.72 Gb for A549 cells (N50 = 1.604 kb). The sequencing reads were aligned to the assembled contigs of all expressed TEs predicted based on our short-read RNA-seq data using minimap2 with the ‘-ax splice -uf -k14’ options, specifically used in spliced long read alignment. SAMtools was used to sort and index the BAM files. bedtools was used to transform BAM files to BED files. The expressed TEs were verified in cell lines, as there were reads that aligned to both the inserted regions and the adjacent exons of the corresponding genes of TEs.

### RNA Extraction and Retrotransposition PCR for TE Expression

1. RNA extraction and sequencing RNA extraction, cDNA synthesis, and retrotransposition-PCR were performed following the manufacturer’s protocol as previously described [94]. For TE retrotransposition detection, the primer design principles were as follows: considering the uncertain length of the expressed TE, if the TE insertion site was between two exons, the primers were located on the exon, and the length of the PCR product was less than 300 bp, excluding the length of TE. For cases where the TE was inserted in the head or tail of the gene, one primer had to be located in the TE, which more than 100 bp away from the end of the TE, and the final product was less than 500 bp. If a gene contained two TE insertions, two pairs of primers were designed accordingly. The designed primers covered all transcripts to the extent that this was possible. The sequences of the primers are shown in Table S6B. To confirm that the bands detected in the PCR assay were the expressed TE as predicted, we purified (Invitrogen) and sequenced the PCR products on the QNome 3841 machine (Qitan Inc., Chengdu, China). All experiments were performed in triplicate.
2. Data analysis For each sample, the contig sequence containing the expressed TE and the associated gene was assembled using the local assembly strategy described previously. Then, sequencing reads were aligned to the assembled contig using minimap2 with the ‘-ax splice’ options, specifically used in spliced long read alignment. The sorted and indexed BAM files were generated with SAMtools. Finally, the Python module pysam was used to extract reads (only primary alignment) carrying expressed TE sequences and spanning the somatic TE and exons of its associated gene.

### Statistical Analysis

Statistical tests are described in the figure legends. Further details of these tests are described below. Where stated, Fisher’s exact test was performed using the function ‘‘fisher.test’’ within R. The chi-square test was performed using the function “chisq.test” within R. The Kolmogorov‒Smirnov test was used for nonparametric testing between two groups using the function ‘ks.test’ within R. Additionally, wherever stated, the Wilcoxon rank-sum test was used for nonparametric testing between two groups using the function ‘‘wilcox.test’’ within R. False discovery rate (FDR) correction was performed to nominate statistically significant associations (‘‘p.adjust’’ within R). KOBAS [95] (version 3.0) was used for gene set enrichment analysis.

## Data and code availability

The sequencing data and any original code required to reanalyze the data reported in this paper are available from the lead contact upon request.

## Acknowledgments

We thank Liyun Bi from the Center of Precision Medicine, West China Hospital of Sichuan University, Chunyan Yu from the Laboratory of Omics Technology and Bioinformatics, West China Hospital of Sichuan University, Xue Xiao and Chengpin Li from the West China Hospital of Sichuan University for their technical assistance. We thank Wei Huang from Northeast Normal University, Kailing Tu and other members of the Dan Xie laboratory for useful discussions. This work was supported by grants from the National Natural Science Foundation of China (92159302 to W. Li, 82173383 to D. Xie, 32170592 and 32370628 to Z. Wang, 3220050 to L. Xia), the Sichuan Province Science and Technology Program (2022NSFSC1553 to L. Xia), and the 1·3·5 project for disciplines of excellence, West China Hospital, Sichuan University (ZYYC20006 to D. Xie). We used external LUSC SRS datasets (ICGC LUSC-KR project) with support from the International Cancer Genome Consortium (ICGC).

## Author contributions

A. X., W.Z, T.Z, X.P. and H.W. contributed equally to this work. They were responsible for the experimental design, execution, data analysis, and manuscript preparation. G.Z., Y.L. and Y.Q. were responsible for sequencing and data cu-ration. X.W., X.L. were responsible for formal analysis and methodology. Y.D. and F.Z. were responsible for validation. D.X. and W.L. were equally responsible for supervision of research, data interpretation and manuscript preparation.

## Competing interests

Dan Xie is a co-founder and shareholder of QitanTech.

## Notes

### Competing Interest Statement

Dan Xie is the co-founder and shareholder of QitanTech.

